# Pre-clinical evaluation of a gene therapy candidate for *SOD1*-ALS shows improved survival and signs of inflammation in the CNS of treated mice

**DOI:** 10.1101/2025.10.16.682744

**Authors:** Pezet Sonia, Hua Jennifer, Marais Thibaut, Delamare Marine, Elouej Sahar, Lemos P. Julia, Castiglione Alexia, Astord Stéphanie, Cohen-Tannoudji Mathilde, Georges Arielle Peche, Rigamonti Mara, Daniele Nathalie, Genries-Ferrand Sandrine, Buscara Laurine, Ratti Antonia, Bohl Delphine, Smeriglio Piera, Biferi Maria Grazia

## Abstract

Amyotrophic lateral sclerosis (ALS) is a progressive neurodegenerative disorder characterized by motor neurons loss (MN). In 15–20% of familial ALS cases, mutations in the superoxide dismutase 1 (*SOD1*) gene are the underlying cause. Targeting human SOD1 (hSOD1) toxicity has emerged as a promising approach to treat SOD1-ALS. We previously demonstrated the efficacy of an exon-skipping strategy using a self-complementary AAVrh10-U7-hSOD1 vector in SOD1^G93A^ mice achieving significant hSOD1 silencing. In this study, we optimized the therapeutic protocol by conducting a dose-finding and biodistribution study of scAAVrh10-U7-hSOD1 following a single intracerebroventricular injection in adult SOD1^G93A^ mice. Our findings demonstrate a dose-dependent reduction in mutant hSOD1 levels in the cortex, spinal cord, and peripheral tissues, sustained for up to 60 days post-injection. *In vivo*, some adverse effects were noted mostly at the highest dose, with inflammation early post-injection and persistent microglial activation in the brain observed around the injection site. Importantly, the medium-dose treatment extended mean survival by up to 27% with a much milder early toxicity, which will provide a great possibility for future applications. Additionally, no major off-target effects were observed in human cell models, highlighting the targeting specificity of this approach and the potential safety for translation. These findings confirm and extend the therapeutic potential of scAAVrh10-U7-hSOD1 gene therapy while emphasizing the need for further technological development to minimize adverse effects and maximize potential clinical benefit.

## INTRODUCTION

Amyotrophic lateral sclerosis is a devastating, fast-progressing disorder characterized by the degeneration of motoneurons in the brain and the spinal cord. These events lead to progressive muscle atrophy, paralysis and ultimately death, typically occurring 3 to 5 years after diagnosis (1). It is estimated that approximately 90% of ALS cases are sporadic while the remaining 10% are familial forms (fALS). Among these inherited forms, 15-20% are caused by mutations in the superoxide dismutase 1 (*SOD1*) gene (2), with nearly 200 mutations identified to date (3), including the G93A mutation reproduced in the most widely studied mouse model of fALS (SOD1^G93A^) (4).

Although the mechanisms underlying SOD1 toxicity are not yet fully elucidated, there is substantial evidence supporting a toxic gain of function of the autosomal-dominant mutations in *hSOD1* (5). The primary deleterious mechanisms associated with the mutated SOD1 involve conformational instability, leading to aberrant misfolding and the formation of toxic aggregates. Given that these defects are considered major drivers of the disease, two primary therapeutic interventions are being explored as treatment options for SOD1-ALS: 1) targeting misfolded SOD1 protein and 2) silencing mutated *hSOD1* mRNA.

*1)* One approach for the first category involves the removal of SOD1 aggregates through the delivery of monoclonal antibodies (6,7) and has been suggested as a more promising approach to treat animal models of SOD1-ALS than the previously tested immunization strategies (8,9). Alternatively, targeting proteostasis through the administration of Sephin 1, a holophosphatase inhibitor, has been shown to prevent motor, morphological, and molecular defects in the SOD1-ALS mouse model by reducing the accumulation of insoluble mutant SOD1 (10,11).
*2)* For the mRNA silencing approach, the recent US Food and Drug Administration (FDA) approval of Tofersen, an antisense oligonucleotide (ASO) that reduces hSOD1 synthesis, represents a major breakthrough in the treatment of SOD1-ALS. Clinical trials have shown promising results (12–14), with Tofersen treatment leading to a reduction in plasma neurofilament light chain (NfL) levels and stabilization of the motor function in few patients, as measured by the ALS functional rating scale revised (ALSFRS-R) (15–17). However, adverse effects including pleocytosis and intrathecal immunoglobulin synthesis in the cerebrospinal fluid (CSF) suggesting an autoimmune inflammation of the central nervous system, have been reported in some patients (18). Additionally, the intrathecal injection protocol is invasive and requires repeated doses, a burden for the patients. Several other methods can be employed to modulate gene expression *in vivo* including RNA interference (RNAi) approaches and CRISPR/Cas9 gene editing (19,20). Recently, uniQure’s AMT-162, an adeno-associated viral vector (AAV) gene therapy candidate expressing a microRNA that binds to *SOD1* mRNA, was administered intrathecally to the first patient in a phase 1/2 clinical trial (https://uniqure.gcs-web.com/news-releases/news-release-details/uniqure-announces-dosing-first-patient-phase-iii-clinical-trial) (21–23). Furthermore, following recent advancements in CRISPR/Cas9 technology, various attempts to edit the *SOD1* gene have begun to emerge (24).

We previously reported the efficacy of gene therapy-mediated exon-skipping strategy against mutant *hSOD1* in the SOD1^G93A^-mouse model (25). This strategy relies on the use of a self-complementary (sc)-AAV vector, which ensures stable and long-term transgene expression with a single delivery. Specifically, we utilized the AAVrh10 serotype, which demonstrated effective central nervous system (CNS) transduction (26), targeting multiple cell types in the spinal cord, including MN (27). This serotype has also been shown to have a low immunogenic profile and to be safe when delivered to the CNS in non-human primates (NHP) (23,28,29). To induce *hSOD1* silencing, the viral vector expresses antisense sequences embedded in the optimized small nuclear RNA U7. Combined intravenous (i.v) and intracerebroventricular (i.c.v) injection of scAAVrh10-U7-hSOD1 in the SOD1^G93A^ mice resulted in a significant reduction of hSOD1 levels in the central nervous system (CNS) and peripheral organs, inducing an extension of survival of up to 92% and 58% in mice injected either at birth or at 50 days of age, respectively (25).

To advance the pre-clinical development of this promising approach, some technical issues needed to be addressed: the high vector dose delivered to the animals (4.5 x 10^14^ vg/kg) and the use of the combined route of administration (ROA).

In this study, we present the results of a dose finding study and a biodistribution analysis of scAAVrh10-U7-hSOD1 in SOD1^G93A^ mice. We also thoroughly assessed the safety profile of the therapy, including the evaluation of potential off-targets effects *ex vivo* and characterization of the early *in vivo* response to gene therapy.

First, we did not detect any major off-target effects following AAV transduction of human cell models *in vitro*, with only a few differentially expressed genes between U7-hSOD1 treatment and controls, indicating a low risk associated with further development of the therapeutic sequence. We then tested three doses after a single i.c.v. injection of the vector in adult SOD1^G93A^ mice, with the highest dose set at 2.4 x 10^14^ vg/kg. Our results showed that a single i.c.v. injection of scAAVrh10-U7-hSOD1 resulted in a significant reduction of mutant hSOD1 levels in the brain cortex and spinal cord, as well as in the periphery as early as 20 days after injection. This downregulation was sustained long-term (up to 60 days post-injection) and occurred in a dose-dependent manner. Furthermore, we observed a maximum of 27% increase in mean survival of mice treated with the medium dose and a delay in disease onset characterized by weight maintenance and preserved motor function, compared to control mice. However, a few animals died 20 days after injection, primarily at the highest doses, and showed microglial activation in the brain at both 20- and 60-days post-injection. These findings are likely due to the procedure of delivering high vector doses in brain’s lateral ventricles.

This study provides a pre-clinical evaluation of the scAAVrh10-U7-hSOD1 candidate and assesses potential safety outcomes associated with AAV-based gene therapy in a rodent model of ALS.

## RESULTS

### Self-complementary AAVrh10-U7-hSOD1 treatment does not induce major off-targets effects in patient-derived cells

As an initial step for the preclinical development of the scAAVrh10-U7-hSOD1 gene therapy, we assessed the potential non-specific targets of the U7-SOD1 construct on patient-derived cells using two relevant models for ALS. In ALS, MN are the primarily affected cells and other cell types, such as skeletal muscle, also contribute to the exacerbation of the pathological phenotype (30). To model this, we used MN differentiated from induced-pluripotent stem cells (iPSC), using a previously published protocol (31) and myoblasts transdifferentiated from fibroblasts via MYOD transduction (32). Each cell type was transduced with either the scAAVrh10-U7-hSOD1 vector or the control vector (U7-CTRL) and we analysed their bulk transcriptomes using RNA-sequencing.

First, we validated the differentiation of iPSC-MN harboring another hSOD1 mutation, by neurofilament (SMI32) staining (Figure 1A). Transduction of these cells with the therapeutic sequence led to a significant and selective downregulation of SOD1 expression. Importantly, we did not observe any off-target gene expression changes compared to control-treated cells in this relevant cell model (Figure 1B).

**Figure 1.**
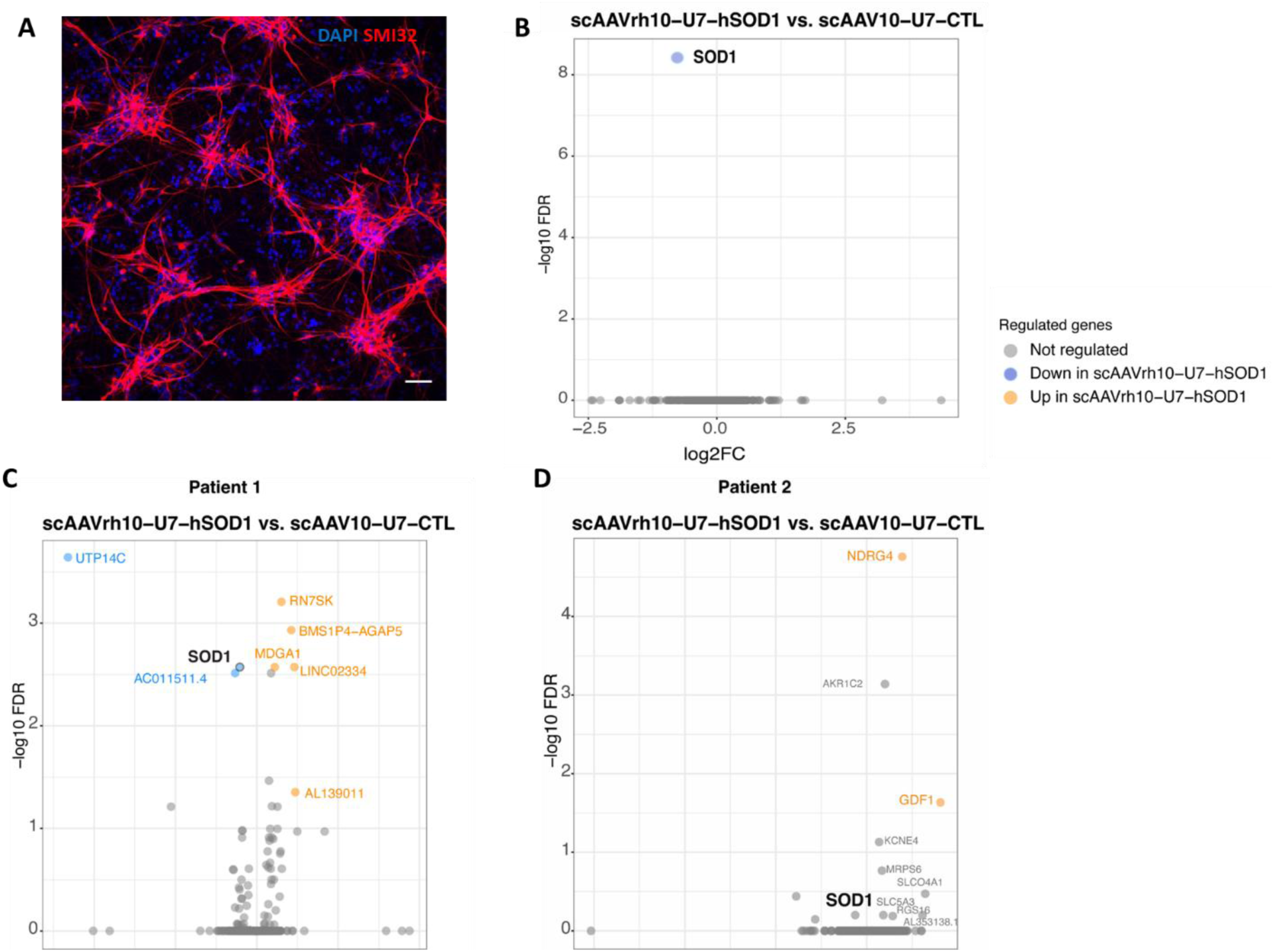
No off-target effects were observed in patients derived cells upon scAAVrh10-U7-hSOD1 treatment. **A.** SMI32 (red) immunostaining labelling of MN-differentiated from induced-pluripotent stem cells from SOD1-ALS patient carrying the N139D mutations at day 25 of differentiation. Scale bar 20µM **B.** Volcano plot representing differentially expressed genes from bulk RNA-sequencing analysis of human MN-differentiated from induced-pluripotent stem cells. LogFC= 1.5 and p-value=0.05. **C, D.** Volcano plot representing differentially expressed genes from bulk RNA-sequencing analysis of human transdifferentiated myoblast from patients harboring a G93D (**C**) or L144F (**D**) mutations. DEG are obtained from the comparison between scAAVrh10-U7-hSOD1 scAAVrh10-U7-CTL treated cells (n=3 independent transduction experiment per group).

Next, we assessed the impact of our viral vector system on myoblasts transdifferentiated from fibroblasts obtained from two patients with different mutations in *hSOD1*. In both cases, the therapeutic sequence did not result in significant alterations in gene expression. For Patient 1, only 8 genes were differentially expressed (Figure 1C), while for Patient 2 only 2 genes were significantly upregulated (Figure 1D). These genes were mainly related to mitosis, which may be attributed to a transdifferentiation batch effect. Importantly, none of these genes were associated with significantly altered biological pathways, suggesting that the therapy did not induce major off-target effects. Additionally, SOD1 expression was reduced by approximately two-fold in Patient 1’s myoblasts following treatment, although this downregulation was not statistically significant in Patient 2. The results demonstrate effective *in vitro* targeting of the mutated hSOD1 with no major off-target effects in patient-derived transdifferentiated myoblasts.

Together, these data suggest that the viral particle expressing the U7-SOD1 construct does not induce major changes in the transcriptome of cells obtained from ALS patients.

### Combined i.v./i.c.v. and single i.c.v. administration of scAAVrh10-U7-hSOD1 outperform i.v. delivery in adult SOD1^G93A^ mice

We previously demonstrated the efficacy of scAAVrh10-U7-hSOD1 gene therapy following combined intracerebroventricular (i.c.v.) and systemic intravenous (i.v.) delivery in SOD1^G93A^ mice. While this approach was promising in term of efficacy, the simultaneous use of these two ROAs could pose safety risks due to high vector exposure to the peripheral tissues, and it may raise regulatory concerns for clinical translation.

To assess whether a single delivery route would be effective in the SOD1^G93A^ mouse model, we conducted a small pilot study comparing the i.v. to the i.c.v. administration, and to the combination of the two ROAs. In our original study, the total dose delivered per animal was of 4.5 x 10^14^ vg/kg, with 4.2 x 10^14^ vg/kg given systemically and 3 x 10^13^ given by i.c.v. We aimed to reduce the overall dose in this follow-up study.

SOD1^G93A^ mice (n=6) were injected at postnatal day 50 (P50), corresponding to an early disease stage, when muscle denervation is already present (33). The following dosing regimens were tested: 1) i.v. injection of 3 x 10^13^vg/kg, 2) i.c.v. injection of 3 x 10^13^vg/kg or at 5 x 10^13^vg/kg, and 3) combination of i.c.v. and i.v. injection at two doses: i.v. 1.5 x 10^13^vg/kg + i.c.v. 1.5 x 10^13^vg/kg and i.v. 3 x 10^13^vg/kg + i.c.v 3 x 10^13^vg/kg.

All treatment groups showed an increased survival compared to non-injected (NI) mice. Statistically significant extensions in median lifespan were observed in the following groups: i.v. injection at 3 x 10^13^vg/kg of (139 days vs 129 days, p=0.0102, 9% mean survival increase), i.c.v. injection at 3 x 10^13^vg/kg (148.5 days vs 129 days, p=0.0041, 12.9% mean survival increase), i.c.v. injection at 5 x 10^13^vg/kg (159.5 days vs 129 days, p=0.0007, 19.5% mean survival increase) and the co-injection group at 6 x 10^13^vg/kg (153.5 days vs 129 days, p=0.0007, 19.9% mean survival increase) (Figure S1A).

The most significant survival extension was observed in the scAAVrh10-U7-hSOD1 i.c.v-injected and co-i.c.v./i.v.-injected group at 5-6 x 10^13^vg/kg with 19.5% and 19.9% increase in median survival compared to NI, respectively (Figure S1B).

Given the comparable survival benefits observed with i.c.v. and co-i.c.v./i.v. injections, we selected the single i.c.v administration as the more suitable ROA for potential clinical translation.

### A single i.c.v. administration of three different doses of scAAVrh10-U7-hSOD1 extend survival, delay disease onset and rescue disease phenotype in SOD1^G93A^ mice

Based on these results from the pilot study and the absence of in-life adverse events (25), we initiated a dose escalation study, starting with a dose of 6 x 10^13^vg/kg and including two additional higher doses. We injected SOD1^G93A^ adult mice at P50, with scAAVrh10-U7-hSOD1 at 6 x 10^13^vg/kg (low dose), 1.2 x 10^14^vg/kg (medium dose) or 2.4 x 10^14^vg/kg (high dose), all administered unilaterally in the brain’s lateral ventricle (i.c.v.) (Figure 2A). The highest dose in this study was approximately half the total dose administered in our previous pre-clinical test and about a 10-fold increase compared to the dose delivered via i.c.v. injection alone (25).

**Figure 2.**
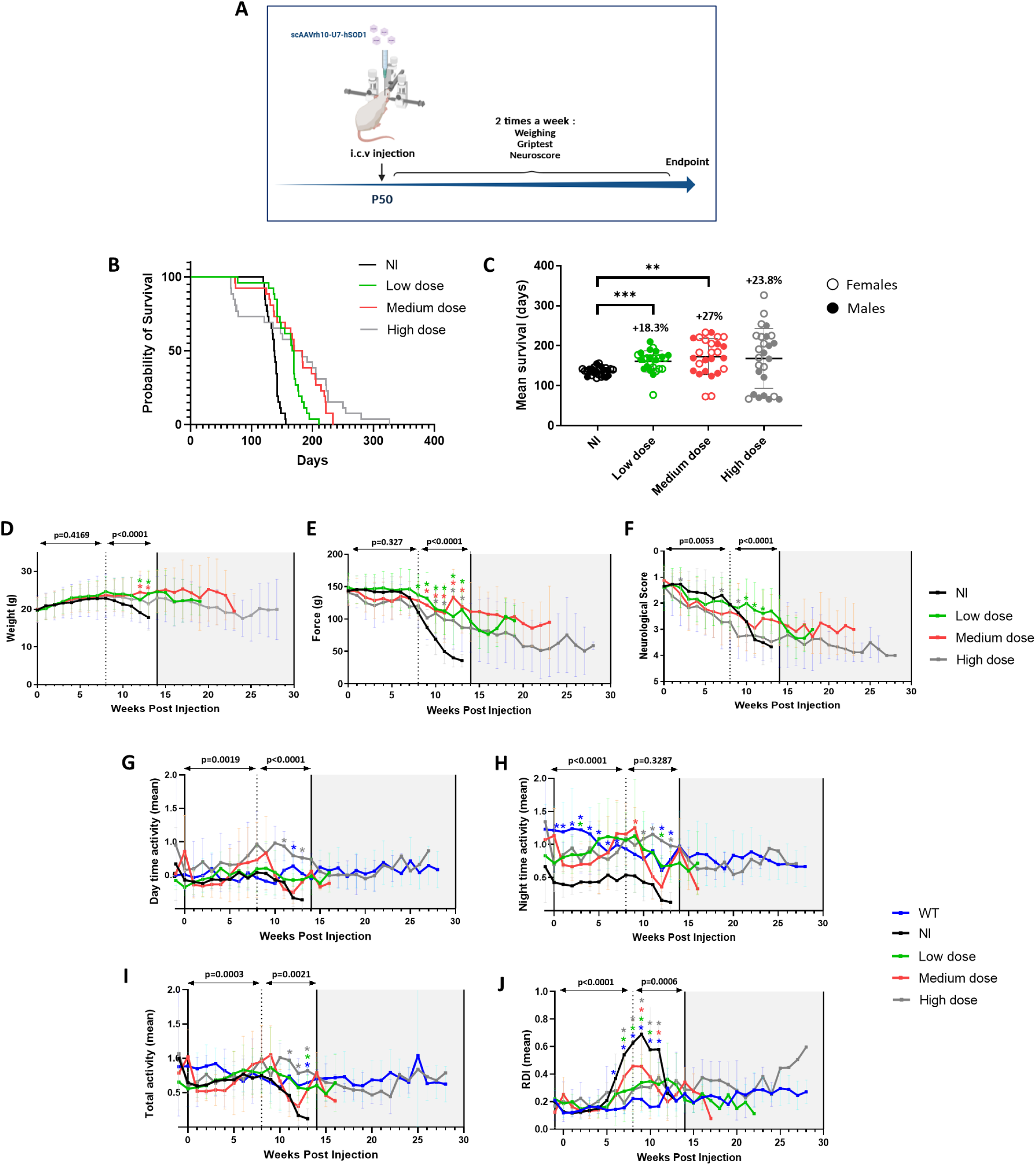
i.c.v scAAVrh10-U7-hSOD1 administration extends survival, delays disease onset and rescues disease phenotype. **A.** Schematic representation of the study design. P50 SOD1^G93A^ mice (n=26, sex-balanced) were injected in the right brain lateral ventricle (i.c.v.) with scAAVrh10-U7-hSOD1 at 6^E^13vg/Kg (low dose), 1.2^E^14 vg/Kg (medium dose) or 2.4^E^14 vg/Kg (high dose). **B.** Kaplan-Meier survival curves of SOD1^G93A^ injected mice and NI mice. Differences between the curves were analyzed using the log rank Mantel-Cox test. ***p = 0.0002; ****p < 0.0001; ****p < 0.0001. **C.** Scatter plot representation of mean survival of scAAVrh10-U7-hSOD1 injected mice compared to NI mice. Data are expressed as the mean ± SD, and differences between groups were analyzed by one-way ANOVA Dunnett’s T3 multiple comparison test, indicating significant differences between the mean survival of NI mice and either scAAVrh10-U7-hSOD1 injected mice at the low dose or at the medium dose (136 ± 10.3 days versus 160.9 ± 26.3 days and 172.7 ± 44.2 days, respectively). **p < 0.01, ***p < 0.001. All percentages of increased survival are compared to the mean survival of NI mice. **D.** Body weight curves, **E.** grip strength curves and **F.** neurological score curves of scAAVrh10-U7-hSOD1-injected SOD1^G93A^ mice compared to NI SOD1^G93A^ mice. *p < 0.05, **p < 0.01, ***p < 0.001, ****p < 0.0001 **G.** Day time activity curves, **H.** Night time activity curves, **I.** Total activity curves and **J.** Regularity Disturbance Index (RDI) curves of scAAVrh10-U7-hSOD1-injected SOD1^G93A^ mice compared to NI SOD1^G93A^ mice. For D to J, differences between groups were analyzed by two-way ANOVA Tukey’s multiple comparison test. All significances (*) are compared to NI group and exact p-values are shown in supplementary Table 1.

To minimize vector batch variability, we used large-scale vector production (see methods) and a large cohort of mice (n=26 per group, sex balanced) for a robust pre-clinical evaluation.

We observed survival extension across all three tested doses, with an 18.3% increase in mean survival for the low dose (160.9 ± 26.30 days versus 136 ± 10.28 days; p <0.001), a 27% increase for the medium dose (172.7 ± 45.18 days versus 136 ± 10.28 days; p <0.01), and a 23.8% increase for the high dose (168.3 ± 74.58 days versus 136 ± 10.28 days) (n=26 per group) (Figures 2B, 2C). Interestingly, the high dose did not lead to a greater survival extension than the medium dose, suggesting that the higher dose was not well tolerated.

In addition to the survival analysis, we performed a comprehensive phenotypic characterization of the treated mice compared to non-injected SOD1^G93A^ mice. All treatment groups exhibited significantly delayed disease onset, as well as phenotypic improvements starting at 8 weeks post-injection. Weight loss was significantly prevented in treated mice throughout the lifespan following vector delivery (p<0.0001) (Figure 2D). Neuromuscular function, assessed using a grip strength test, was preserved in treated mice, while control mice began to show decline. Treated mice exhibited significantly greater strength compared to NI mice (p<0.0001) (Figure 2E). Hindlimb function, assessed by neurological scoring based on hindlimb extension, was significantly improved in the low dose group compared to NI mice (Figure 2F), and this improvement was maintained in treated mice with extended survival.

We further analyzed spontaneous motor activity using a digital system for continuous in-life monitoring of multiple parameters (Digital Ventilated Cages, DVC system). Over a period of up to 28 weeks post-injection, treated mice demonstrated an overall rescue of motor activity. Starting at 8 weeks post-injection, recorded movements were significantly greater in treated mice compared to NI mice, particularly in the high dose, with activity levels comparable to WT mice (p<0.005) (Figures 2G-I). As expected, activity was notably higher during night time, which coincides with the peak activity levels in mice (Figures 2H). Interestingly, the Regularity Disruption Index (RDI)(34), a digital biomarker quantifying irregularities of activity pattern and rest/sleep-related disturbances, was significantly restored in treated mice compared to NI mice, and it was similar to WT animals (p=0.0006). In NI SOD1 mice, RDI increased rapidly starting at 5 weeks, reflecting irregularities in activity patterns (Figure 2J) before paralysis onset, which is consistent with sleep disturbances observed in ALS patients (35).

As we previously reported (25), while half of the treated animals died after developing progressive signs of limb paralysis, 35% of animals in the low dose group, 46% in medium dose group and 77% in the high dose group exhibited atypical ALS signs in the end-stage disease. These animals showed progressive weight loss and decreased limb strength in the final days before reaching the end-point, but they maintained ambulation and a relatively active actimetry profile in the weeks prior. This phenomenon was also described by Stoica et al. (36) in SOD1 mice treated with a microRNA, and it may result from the extended lifespan and modified disease progression due to treatment.

In conclusion, these analyses demonstrate that the treatment induced survival extension and phenotype improvement in SOD1^G93A^ mice. However, we cannot overlook the fact that despite the efficiency of the treatment, early unexpected deaths occurred among treated mice, approximately 20 days post-injection, in a dose-dependent manner (1 early death in the low-dose group, 2 in the medium-dose group, and 7 in the high-dose group, out of 26 mice per group) (Figure 2B). These unexpected events led to further investigations in order to better understand the benefit and risks of this gene therapy.

### i.c.v. gene therapy administration leads to dose-dependent decrease of hSOD1 levels in the CNS and the periphery of SOD1^G93A^ mice

We first completed the pre-clinical characterization of the dose study, by assessing target engagement and biodistribution following administration of the three different doses of scAAVrh10-U7-hSOD1. An independent cohort of SOD1^G93A^ mice (n=4 per group, sex balanced) was injected following the same protocol as the survival study, and tissues were collected at P110, corresponding to end-stage disease in NI mice (Figure 3A) approximately two months after vector injection.

**Figure 3.**
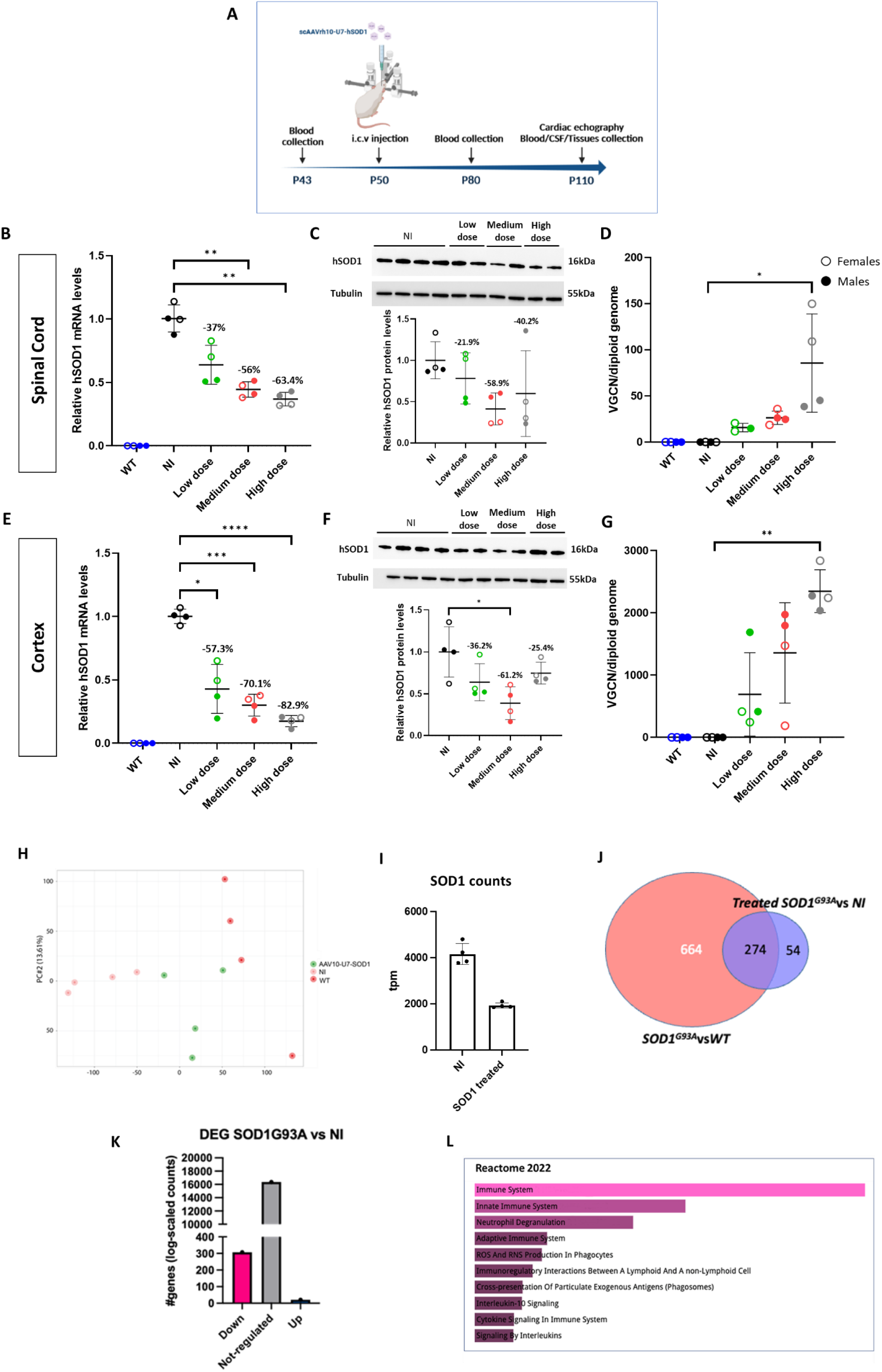
scAAVrh10-U7-hSOD1 administration via i.c.v. results in a dose-dependent decrease of hSOD1 levels in the CNS. **A.** Schematic representation of the study design. P50 SOD1^G93A^ mice (n=4, sex-balanced groups) were injected with scAAVrh10-U7-hSOD1 at 6^E^13vg/Kg (low dose), 1.2^E^14 vg/Kg (medium dose) or 2.4^E^14 vg/Kg (high dose) in the right lateral ventricle (i.c.v.) and tissues were collected at P110. **B, E.** qRT-PCR analysis of *hSOD1 mRNA* levels normalized to *HPRT* and expressed as relative to levels in spinal cord (B) and cortex (E) of NI mice. **C, F, I.** Relative hSOD1 protein levels measured by Western blot and normalized to TUBULIN (spinal cord – **C** and cortex – **F**) and expressed as relative to levels in NI mice. **D, G.** Vector genome copy number (VGCN) per diploid cell in the spinal cord (**D**) and cortex (**G**). Data are expressed as the mean ± SD, and differences between groups were analyzed by one-way ANOVA. *p < 0.05, ** p < 0.01, ***p < 0.001. All percentages are calculated as relative to the group of NI mice. Significances with WT group are not shown on graph. **H.** Principal component analysis plot obtained analyzing the variably expressed genes in each RNA-sequencing sample from spinal cord of SOD1^G93A^ mice treated with the high dose, NI SOD1 ^G93A^ and WT. **I.** Transcript per million (TPM) counts for the *hSOD1* mRNA in gene therapy-injected SOD1^G93A^ mice vs NI. **J.** Venn diagram overlap between the DEG of SOD1^G93A^ spinal cord vs WT spinal cord and the treated SOD1^G93A^ spinal cord compared to the untreated group. **K.** Histogram representing the number of genes that are downregulated (pink bar), upregulated (blue bar) and unchanged (grey bar) in treated SOD1^G93A^ compared to NI SOD1^G93A^ spinal cord. n=4 per group, sex-balanced. **L.** Pathway analysis -Reactome 2022 gene-set collection-of all DEG in treated SOD1^G93A^ compared to NI spinal cord.

We evaluated the reduction in hSOD1 mRNA and protein levels across various organs following therapy administration. A dose-dependent effect was observed in nearly all analyzed compartments, including the spinal cord, brain, skeletal muscle and peripheral organs. The low dose was already sufficient to induce a 37% reduction of *hSOD1* mRNA in the spinal cord (Figure 3B) and 57.3% decrease in the brain cortex (Figure 3E). At the highest dose, *hSOD1* mRNA levels were reduced by 65% in the spinal cord (Figure 3B) and 83% in the cortex (Figure 3E). Significant reductions in hSOD1 protein levels were also noted in the spinal cord and brain (Figure 3 C, F). Significant reduction of hSOD1 mRNA was also observed in other brain regions, heart and liver (Figure S2A).

In parallel with the observed target engagement, the vector genome copy number (VGCN) increased with the administered doses (Figures 3D, G). The cortex showed high transduction, likely due to its proximity to the injection site, whereas lower transduction was detected in skeletal muscles, notably triceps, tibialis anterior and gastrocnemius (Figure S2B), as expected with an i.c.v. injection in adult mice.

In muscle, the low-dose injection did not result in a reduction of *hSOD1* mRNA levels (Figure S2A) whereas the high dose led to a greater knockdown, which was approximately double compared to the medium dose in the triceps (Figure S2A). Similar results were obtained in the gastrocnemius muscle, though no reduction in *hSOD1* mRNA was observed in the tibialis anterior, suggesting variable effects in different skeletal muscles types (Figure S2A).

A modest downregulation of hSOD1, was also observed in serum and cerebrospinal fluid (CSF) of treated mice compared to NI controls (Figure S2C, D), suggesting that analysis of these biofluids could be a potential method for future monitoring of therapeutic effects in patients.

To assess whether our therapeutic approach could induce a molecular rescue in the spinal cord at P110, we performed RNA-sequencing. Principal component analysis (PCA) of spinal cord extracts revealed low variance between SOD1^G93A^ mice treated with the highest dose and WT mice, suggesting effective rescue of pathological features in the spinal cord (Figure 3H). About 45% of variance in PC1 was observed between the diseased SOD1^G93A^ spinal cord and the WT controls (Figures S2G), with a wide range of gene expression differences among these groups (Figure S2E, F).

Gene therapy administration at the highest dose resulted in a 50% decrease in *hSOD1* mRNA expression in the spinal cord of treated mice compared to NI SOD1 mice (Figure 3I). Only 1/3 of the differentially expressed genes (DEG) between the NI diseased mice and the WT controls remained unchanged after treatment (274 out of approximately 1000 DEG) (Figure 3J). Treatment administration induced the downregulation of 300 genes and upregulation of only 21 genes in the spinal cord (Figure 3K). These differences in gene expression were associated with an enrichment in several pathways, including regulation of reactive oxigen species (ROS) (Figure 3L), suggesting that the treatment at the highest dose induced an inflammatory response. In the triceps, we observed a moderate reduction in *hSOD1* expression (1.6 fold, pvalue=0.0072) as expected, given the limited muscle transduction (Figure S2B) with 147 DEGs between injected and non-injected muscles (Figure S2H).

### i.c.v. administration of scAAVrh10-U7-hSOD1 in SOD1^G93A^ mice induces early, transient impairments primarily at high treatment dose

To investigate the cause of the early deaths observed within 2 weeks post-injection (Figure 2A and B), a small cohort of SOD1^G93A^ mice (n=4 per group, sex balanced) was thus injected with scAAVrh10-U7-hSOD1 at P50. Tissues were collected 20 days later, at P70 (Figure 4A), a time point corresponding to the observed deaths.

**Figure 4.**
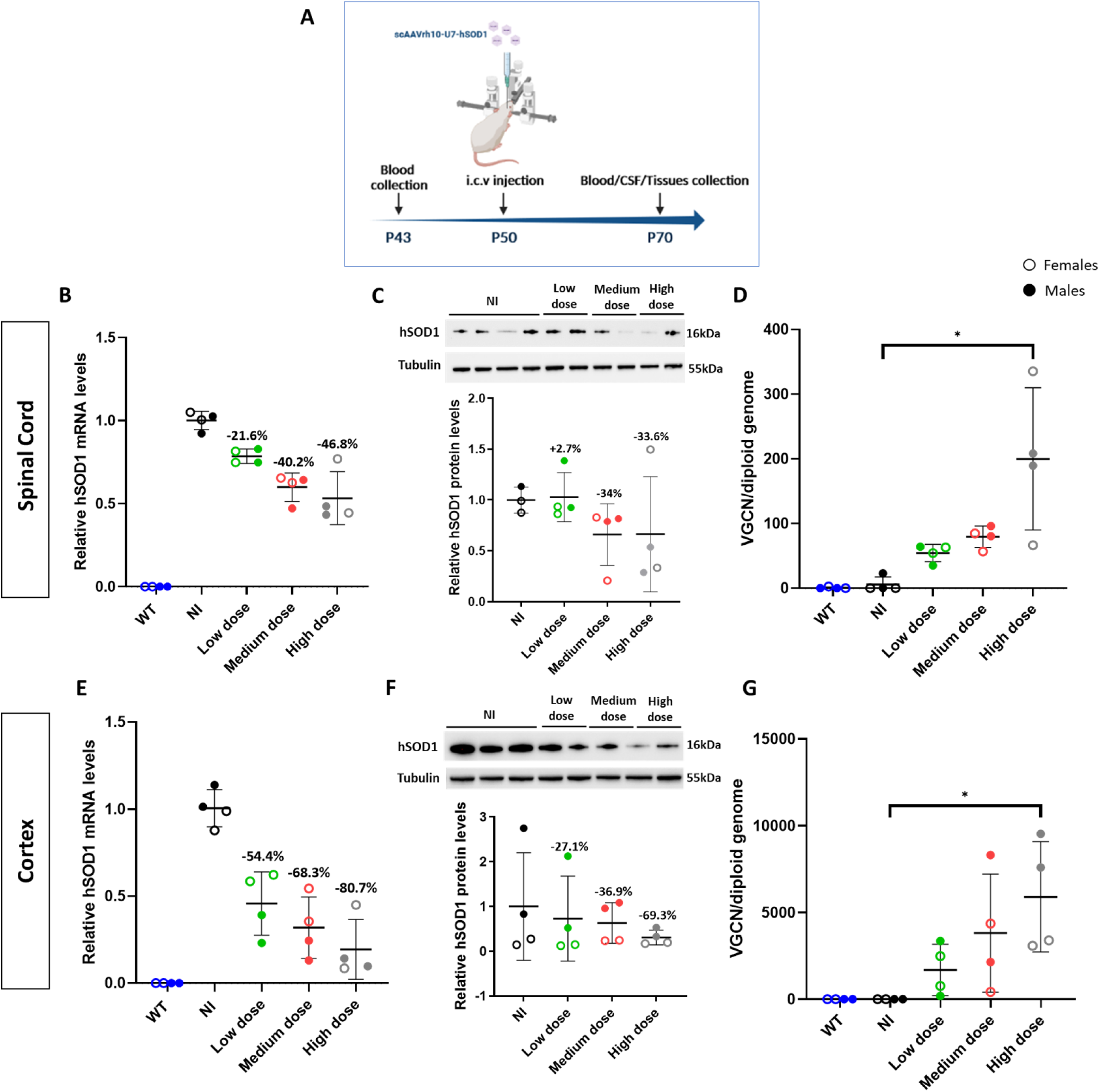
i.c.v scAAVrh10-U7-hSOD1 administration in SOD1^G93A^ mice induces an efficient decrease in hSOD1 levels in the CNS early after injection in a dose-dependent manner. **A.** Schematic representation of the study design. P50 SOD1^G93A^ mice (n=4, sex-balanced groups) were injected with scAAVrh10-U7-hSOD1 at 6^E^13vg/Kg (low dose), 1.2^E^14 vg/Kg (medium dose) or 2.4^E^14 vg/Kg (high dose) in the right lateral ventricle (i.c.v.) and tissues collected at P70. **B, E.** qRT-PCR analysis of hSOD1 mRNA levels normalized to HPRT mRNA and expressed as relative to levels in spinal cord (**B**) and cortex (**E**) from NI mice. **C, F.** Relative hSOD1 protein levels measured by Western blot, quantification of hSOD1 protein levels was normalized to TUBULIN and expressed as relative to levels in NI spinal cord (**C**) and cortex (**F**). **D, G.** Vector genome copy number (VGCN) per diploid cell in spinal cord (**D**) and cortex (**G**). Data are expressed as the mean ± SD, and differences between groups were analyzed by one-way ANOVA. *p < 0.05, ** p < 0.01, ***p < 0.001. All percentages are compared to the group of NI mice. Significances with WT group are not represented.

We first verified whether tissue transduction and corresponding target engagement occurred as early as 20 days post-injection. VGCN analysis revealed the highest viral particle concentration in the CNS of mice treated with the highest dose, compared to NI controls (Figure 4D, G). This finding was consistent with results observed 2 months post-injection in all analyzed tissues, except in cortex, where viral particles were higher, likely due to the injection procedure. The presence of vector genomes correlated with a significant 80% reduction in *hSOD1* mRNA in the cortex and 47% in the spinal cord (Figure 4B, E). Lower doses resulted in a similar significant reduction in *hSOD1* mRNA in the cortex, with a 54% decrease at the lowest dose and a 68% decrease at the intermediate dose. However, the effect was less pronounced in the spinal cord, where hSOD1 mRNA levels were reduced by 21% and 40% in the low and medium doses, respectively. This was accompanied by a dose-dependent decrease in SOD1 protein levels in the spinal cord and brain (Figure 4C, F). Interestingly, the treatment did not induce a downregulation of *hSOD1* in the muscle (Figure S3A), likely due to the insufficient viral particles delivery to the triceps, gastrocnemius and tibialis anterior (Figure S3B). Surprisingly, however, efficient target engagement was observed at this early stage in the serum of treated mice (Figure S3C) but not in the cerebrospinal fluid (Figure S3D).

These analyses confirmed successful transduction and efficient reduction of *hSOD1* levels in the CNS and systemic circulation 20 days after injection.

To further explore potential causes of the observed early deaths, we analyzed the serum chemistry of treated mice at P70 and compared it to P110 samples. We assessed liver function (alanine aminotransferase, aspartate aminotransferase), renal function (creatinine), and cardiac markers (creatinine phosphokinase, blood urea nitrogen) as well as brain, liver and bone markers (alkaline phosphatase) that could putatively correlate with sudden deaths. No significant differences were observed between treated and NI mice, as well as between treated mice and WT controls, at either time point (P70 or P110) (Figure 5A, B).

**Figure 5.**
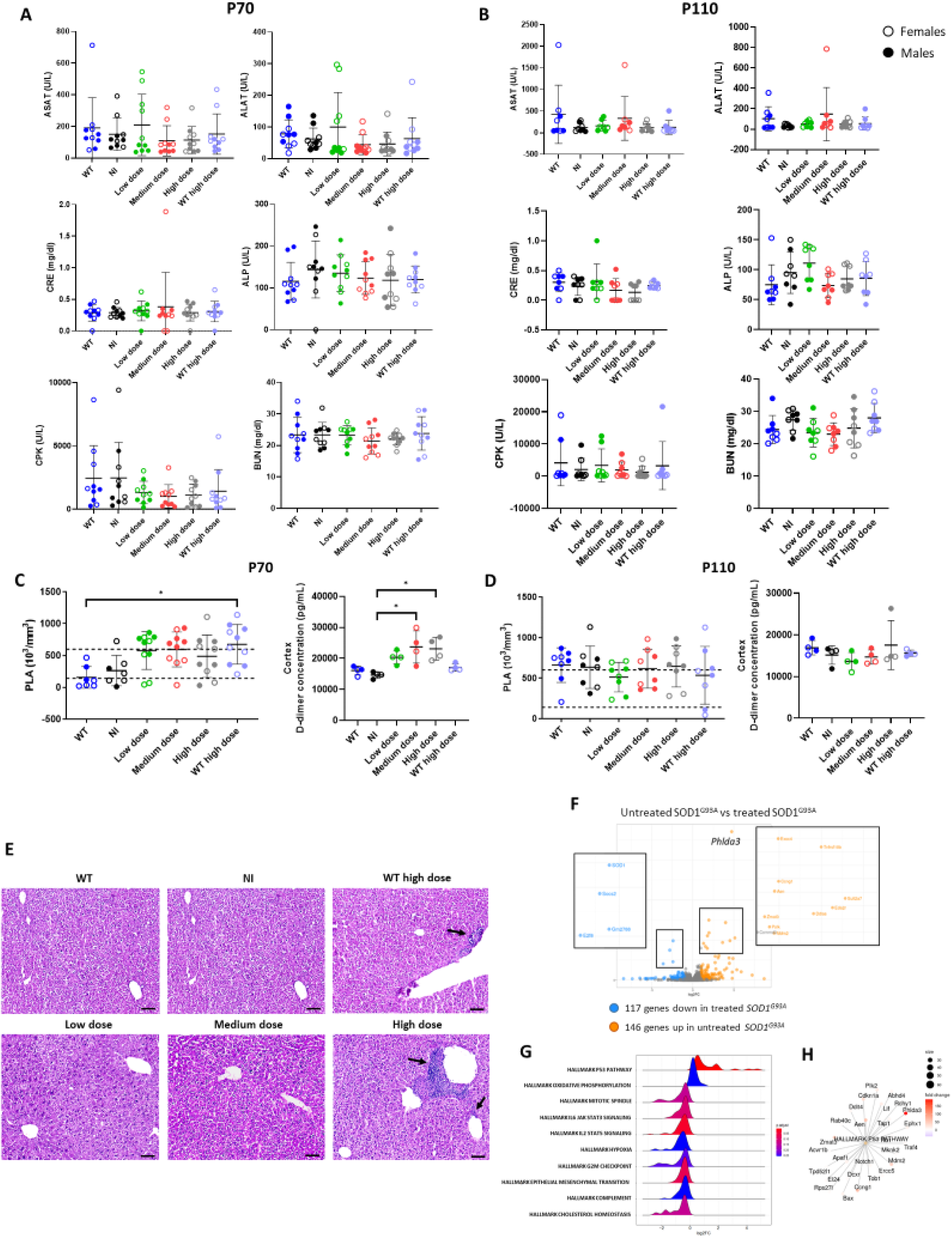
i.c.v scAAVrh10-U7-hSOD1 gene therapy induces early and transient impairments in treated SOD1^G93A^ mice in a dose-dependent manner. **A, B.** Scatter plot representation of serum markers quantification in injected, non-injected SOD1^G93A^ and WT mice (n=10, sex-balanced groups) at P70 (A) or P110 (B); aspartate aminotransferase (ASAT), alanine aminotransferase (ALAT), creatinine (CRE), alkalin phosphatase (ALP), creatinine phosphokinase (CPK) and blood urea nitrogen (BUN). Data are expressed as the mean ± SD, and differences between groups were analyzed by one-way ANOVA. **C, D.** Scatter plot representation of: platelets (PLA) levels (left panels) in peripheral blood of injected, non-injected SOD1^G93A^ and WT mice (n=10, sex-balanced groups) (dotted line represents physiological levels of platelets) and D-dimer concentration (right panels) in the cortex, normalized to total protein concentration (n=4, sex-balanced groups) at P70 (**C**) or P110 (**D**). Data are expressed as the mean ± SD, and differences between groups were analyzed by one-way ANOVA. *p < 0.05. **E.** Hematoxylin & Eosin staining of representative transverse section of liver from injected and non-injected WT and SOD1^G93A^ mice (n=6, sex-balanced groups) of P70 (A). Black arrows show perivascular infiltrates. **F.** Volcano plot representing upregulated and downregulated genes in scAAVrh10-U7-hSOD1 treated SOD1 day 20 post-injection vs untreated SOD1 liver extracts; LogFC= 1.5 and p-value=0.05, 146 genes are upregulated and 117 are downregulated. **G.** Density plots of signaling pathways representing the frequency of log2FC values of up and downregulated genes from liver of SOD1^G93A^ mice treated with the high dose vs untreated. **H.** Gene-network plot showing the upregulated genes for the p53 pathway in treated SOD1^G93A^ liver vs untreated liver.

No statistically significant changes in inflammatory cytokine levels were observed between the groups at early or late time points (Figure S4A), although high variability was noted, likely due to the low sample size.

To ensure that the observed effects were not disease-related, we also injected WT mice with the high dose, and similar heterogenous patterns were observed (Figure 5 A-D).

In the hematology assessment (Figure S4 B, C), we observed an increase in platelet counts in treated mice compared to NI and WT controls at P70, but not at P110 (Figure 5C, D). As an elevated platelet count can lead to blood clot formation and thrombosis, we assessed D-dimer levels, a biomarker of thrombosis previously described in microangiopathy events following AAV gene therapy administration (37, 38). Indeed, a statistically significant increase of D-dimer was detected in the cortex of medium- and high-dose treated mice at P70 compared to controls, suggesting the occurrence of a brain thrombosis event, which may be correlated to the observed sudden early deaths.

In contrast to previous reports (38), the immunological profile of the anti-AAV10 antibodies in the serum of treated mice did not correlate with the thrombosis. In fact, we detected a higher level of antibodies at 2 months after injection than 20 days after injection, when compared to controls (Figure S4D, E). This finding contrasts with the notion that anti-capsids antibodies are the primary cause of thrombotic microangiopathy (38), suggesting that a different, yet-to-be identified mechanism may be involved in our model. Moreover, we did not detect an adaptative T-cell response (Figure S4 F, G), which has been previously associated with secondary effects of gene therapy. FOXP3+/CD25+ Treg cells were not upregulated in the spleen or lymph nodes of treated mice at either early or late time points. In fact, we observed a decrease in Treg levels in treated mice compared to NI mice, with levels closer to those in WT animals (Figure S4F, G). The Treg response is critical for sustained transgene expression and long-term efficacy (39), but it is typically dampened in intra-tissue injections (e.g. liver) compared to systemic administration (40). No significant difference in Treg populations were observed upon treatment at the late time-point.

Despite high rate of transduction of the liver (Figure S3B), no elevation in transaminase levels was detected in treated mice across all doses (Figure 5A, B), suggesting no acute hepatotoxicity. This event has been previously reported in several gene therapy-based clinical trials (41–43) and non-clinical studies (44). Histological analysis of liver tissue showed perivascular inflammatory infiltrates as well as hepatocyte degeneration in one SOD1 mouse and one WT mouse, both injected with the high dose (Figure 5E, black arrows). However, this histopathologic manifestation was also observed in WT mice without treatment and does not represent conclusive evidence of toxicity. Nevertheless, approximately 250 genes were differentially expressed in the liver of high-dose treated SOD1 mice compared to NI controls, with enrichment in apoptosis and oxidative stress pathways (Figure 5F, G, H). These transcriptomic changes, observed early after injection, suggest acute perturbation in liver homeostasis that do not result in long-term toxicity, as shown by the stable *hSOD1* downregulation at P110 (DEG: ∼50 with a consistent 2.5 to 2.7-fold reduction in hSOD1).

Finally, we assessed cardiac health in treated mice, since heart toxicity has also been reported following AAV gene therapy (45, 46). Levels of blood urea nitrogen (BUN), a marker for heart failure, were similar between groups at both downregulation P70 and P110 (Figure 5A, B) as well as the cardiac troponin 3 levels (Figure S4H, I). Histology revealed no structural changes (data not shown), suggesting the lack of acute heart toxicity.

Cardiac echography was assessed only in mice injected at P110 and showed no differences in ejection fraction (EF) and fractional shortening (FS), both markers of heart failure (Figure S4J). We noticed a decrease in end-diastolic volume in treated mice compared to WT mice, suggesting potential cardiac hypertrophy and dilatation due to high blood pressure. However, this change was not significant when compared to NI mice, and may represent an intrinsic characteristic of the SOD^G93A^ mouse model, possibly accentuated by the gene therapy administration.

In conclusion, despite a comprehensive analysis, we identified only a few impairments occurring in treated mice 20 days post-treatment, such as sign of brain thrombosis and transcriptomic changes in liver. Both of these effects were resolved 2 months after injections, but may explain the early deaths observed in the treated groups.

### Microglia activation is induced early after scAAVrh10-U7-hSOD1 administration and persists in the brain of treated animals

High doses of viral vector have been implicated in dorsal root ganglia (DRG) toxicity, as observed in few preclinical trials in pigs and non-human primates, as well as in clinical trials involving SOD1 patients (21, 41, 47, 48). To investigate this potential issue in our model, we assessed the effect of our vector on DRG integrity. However, we observed no evidence of inflammation or cell death in any of the groups and at any time point (Figure S5A), thus excluding DRG toxicity in our model.

To identify other potential short-term detrimental effects of the therapy, we analyzed microglial activation and astrocytes activation in the CNS (brain and spinal cord) at P70. We found increased microglial signal, as quantified by IBA1 staining, in the brain and spinal cord of mice treated with the high dose compared to NI controls. However, astrogliosis, measured by GFAP staining, was similar in the brain of treated SOD1 mice and NI controls (Figure 6A, C).

**Figure 6.**
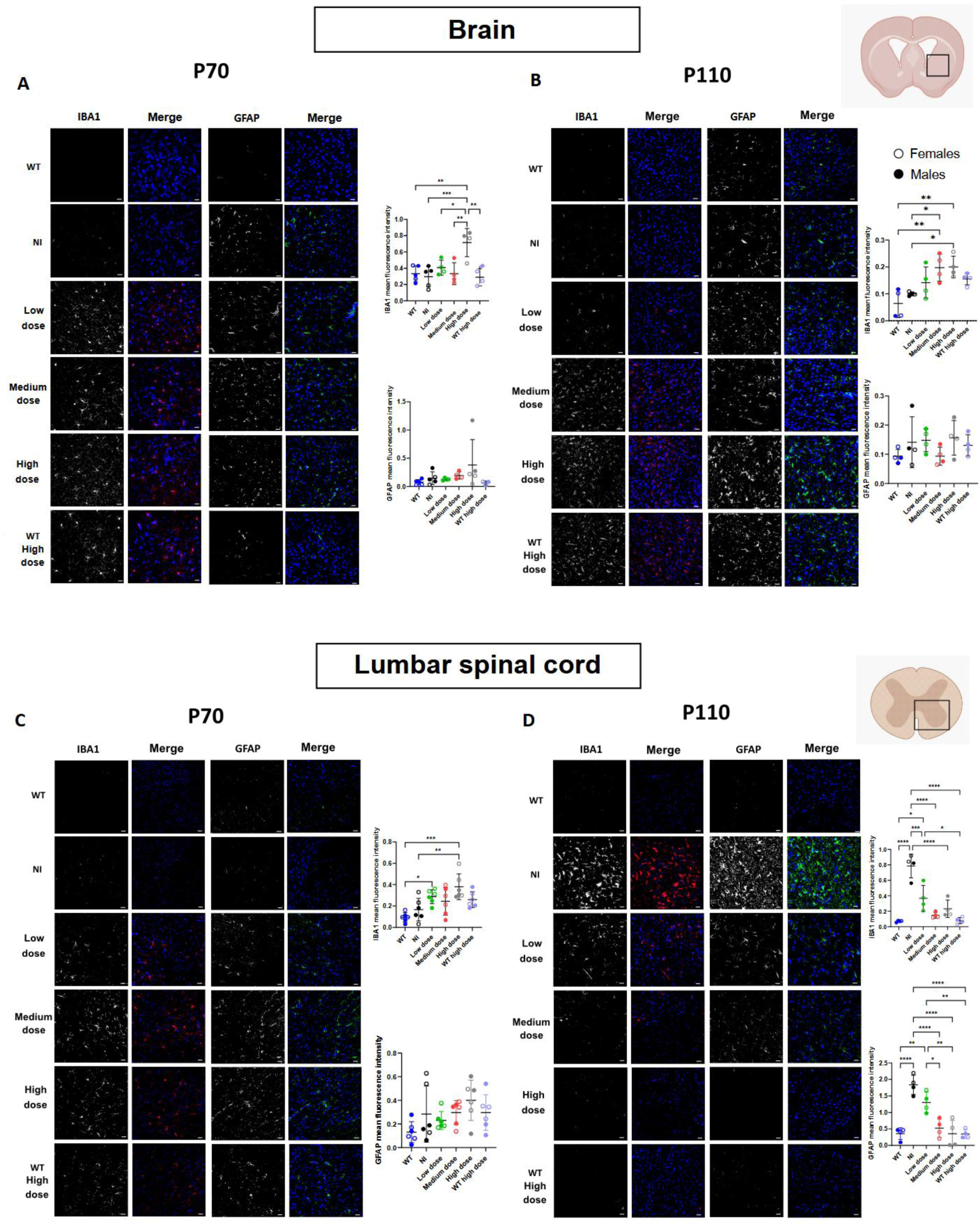
i.c.v scAAVrh10-U7-hSOD1 administration to SOD1^G93A^ mice induces early and transient neurotoxicity in a dose-dependent manner. **A, B.** IBA1 (red) and GFAP (green) immunostaining labelling of representative transverse section of the brain (region as depicted in the brain scheme in the upper right) from injected and non-injected WT and SOD1^G93A^ mice (n=6, sex-balanced groups) of P70 (**A**) and P110 (**B**) and quantitative analysis of mean fluorescence intensity of GFAP and IBA1 immunostaining. **C, D.** IBA1 (red) and GFAP (green) immunostaining of representative transverse sections of the ventral horn of the lumbar spinal cord (region as depicted in the spinal cord scheme in the upper right) from injected and non-injected WT and SOD1^G93A^ mice (n=6, sex-balanced groups) at P70 (**C**) and P110 (**D**) and quantitative analysis of mean fluorescence intensity of GFAP and IBA1 immunostaining. Scale bar 50 µM. Data are expressed as the mean ± SD, and differences between groups were analyzed by one-way ANOVA. *p < 0.05, ** p < 0.01, ***p < 0.001, ****p < 0.0001.

Importantly, microglial and astrocytic activation in the spinal cord was reduced in a dose dependent-manner, consistent with previous findings in SOD1 mouse model treated with U7-based gene therapy (25). In contrast, activation persisted in brain regions adjacent to the ventricles, even two months after vector injection (Figure 6B, D). This result suggests that high-dose treatment may cause long-term brain damage, possibly due to sustained vector presence at the injection site.

Additionally, we observed a dose-dependent enlargement of the lateral ventricle in both SOD1-affected and WT mice two months post-injection (Figure S5 B, C), indicating a physical effect of the procedure on the mouse brain. Enlarged perivascular spaces have been observed in the brain of patients suffering from thrombotic microangiopathy (TMA) (49), which links this finding to the observed increase in D-dimer levels. Based on these observations, we hypothesize that high doses of viral vector in the brain could induce neurotoxicity and a similar microangiopathy-like response in the brain, although we did not detect an increase in creatinine levels typically associated with TMA (Figure 5A, B). It remains possible that treated mice develop TMA following AAV injection without exhibiting all of the symptoms seen in patients.

Taken together, these data represent the first demonstration in a rodent model that high load of gene therapy vectors may induce transient safety concerns. In most animals, these alterations resolve within a few weeks; however, in some cases they may be fatal. Despite the strong potential of this gene therapy in rescuing central nervous systems as well as peripheral abnormal levels of toxic hSOD1, predicting which animals will experience these effects and how many will be affected remains a significant challenge.

## Discussion

This study aimed to develop a gene therapy approach utilizing a U7-AS construct to induce exon 2 skipping in the *hSOD1* gene. Our first experiments evaluated the safety of the therapeutic molecule in two independent ALS patient-derived cellular models, finding no potential off-target effects, as confirmed by transcriptomic analysis following treatment.

We then proceeded with *in vivo* testing. The study design was focused on two key considerations: reducing the previously tested dose and administering a single i.c.v. delivery. This ROA has been safely tested in both pediatric and adult patients with various disorders (50, 51), including ALS (clinical trial: NCT01384162), further supporting its potential for clinical translation.

Our treatment induced long-term target engagement, with an efficient and sustained decrease in hSOD1 levels observed in both the CNS and the periphery of treated mice for up to two months. Additionally, we saw improvements in survival and motor function of SOD1 ALS mice. A transcriptomic analysis revealed that the therapy restored approximately two-thirds of the differentially expressed genes in the spinal cord of treated mice compared to NI animals. However, we noticed enrichment of pathways associated with inflammation and early animal death, particularly in the high-dose treated group. To further explore these safety findings, we characterized these events in detail.

We hypothesized that the unexpected deaths were dose-dependent, with one animal dying in the low-dose group, two in the medium-dose group, and seven in the high-dose group. These fatalities were not observed in our previous study, where we used a total high dose of 4.5 x 10^14^ vg/kg (25). In that study, the dose was primarily delivered systemically, with only 3×10^13^ vg/kg administered via i.c.v. injection. In contrast, in the current study, the lower i.c.v. dose was already double the previous dose, and it is possible that the high vector dose directly introduced into the brain ventricle caused some dysfunctions. This suggests that although i.c.v. injection alone yields better results than i.v. delivery (as shown in this and previous studies)(52–54), a combined ROA may still be beneficial and tolerable, at least in ALS pre-clinical models.

Another factor to consider is that to deliver the three doses in this study, we injected a relatively large volume (1µL per gram of body weight, or approximately 20-30 µL per mouse) directly into the brain lateral ventricle. This could have contributed to the observed enlargement of the brain cavity (Figure S5A). It is possible that removing some of the injected volume from the CSF could help reduce the increased pressure on the ventricle, as is commonly done in intrathecal (IT) antisense administrations (55). However, this approach would need to be tested in mice to determine whether it could offer any benefit to the procedure.

Overall, we hypothesize that the injected dose and volume given by i.c.v. in this study may have been too high for mouse studies and should be reduced in future studies to avoid potential safety concerns. However, this will require further technical optimizations, as the need for a highly concentrated vector titer to fit in the reduced volume for injection in the lateral ventricle represent a significant barrier to translation. In fact, current biomanufacturing pipelines have limited capacity to produce highly concentrated viral therapies in the small volume required for i.c.v. injection in mice. This issue may also apply to human applications, and we anticipate an urgent need for new production methods to achieve better yields while maintaining high vector quality for CNS targeting.

A potential alternative solution to this problem could be to explore other methods for widespread brain and spinal cord transduction, using alternative ROA (56–58) (NCT05866419) or brain penetrating capsids with peripheral de-targeting, which might allow for effective CNS delivery via systemic administration (59, 60).

We believe that the high viral vector doses at the site of injection were responsible for the microglia activation and D-dimer accumulation observed in this study. Despite being an immune-privileged site, the elevated viral vector presence in the brain compartment could have contributed to ventricular enlargement and resident microglia activation. Microglial activation and astrogliosis, both markers of neurotoxicity, were observed 20 days post-injection, suggesting that this response might be a more sensitive indicator of AAV vector-induced toxicity than DRG evaluation in mice. Interestingly, in contrast to previous reports (61–63), we did not observe any signs of DRG toxicity, such as immune infiltration, suggested as a parameter to assess toxic effects from gene therapy administration. While our study was not designed as a comprehensive safety assessment, some of the analyzed parameters, including the increased Iba1-positive cells, indicating glial activation, align with recent studies reporting toxicity following AAV injection, particularly at high doses. Notably, the unexpected negative outcomes observed in these studies include neuroinflammation (28) (64), ocular inflammation (65, 66), hepatotoxicity and DRG toxicity (41, 47, 48). This growing body of evidence suggests that high AAV vector doses may induce toxicity independently of systemic host immune responses to the capsid or the transgene (41).

Chronic microglial activation leads to the release of inflammatory molecules, which can cause neuronal death if the activation persists (67). Similarly, excessive and dysfunctional astrogliosis worsens neurodegeneration in ALS (68), making these cell types valuable therapeutic targets. However, further experiments are needed to optimize their targeting, preserve their beneficial functions, and minimize harmful activation.

Although we excluded systemic immune activation, as indicated by blood parameters, Treg analysis, and anti-capsids antibodies level, we detected some indicators of heart damage in a few injected mice, although average values were not statistically different between NI and treated mice.

Further analysis revealed no significant increase in organ toxicity biomarkers or inflammatory cytokines. However, differences in immune responses between humans and mice should be considered, as murine models may not be the most accurate model to study certain inflammatory responses. Indeed, genetic background and environmental exposure in mice do not fully replicate human conditions. As previously noted with DRG toxicity, murine models may fail to fully predict clinical efficacy and safety (45) and similarly, adverse manifestations in animals may not necessarily translate to humans (69). Thus, the limitations of animal models should be carefully considered when designing preclinical studies for gene therapy.

It is also important to acknowledge that our therapy targets the toxic accumulation of mutant hSOD1, and was tested in a transgenic model in which basal endogenous SOD1 is still present. If the widespread targeting observed in mice can be replicated in translational studies, predicting the potential detrimental effects of a significant reduction in SOD1 expression across tissues will be challenging, particularly in organs that were very efficiently targeted in our study, such as the liver and heart. One potential strategy to address this challenge could be the selective targeting of the mutant allele, as previously described (70).

As evidence of AAV gene therapy-induced toxicity continues to accumulate, it is crucial to develop a comprehensive understanding of potential negative side effects. Future efforts should focus on developing next-generation vectors for ALS that reduce viral load while ensuring efficient and safe targeting of the CNS and MNs. The challenge remains to identify more effective, safer and economically sustainable bioproduction methods that will reduce costs and expand patients access to treatment. Despite current limitations, this study has confirmed the efficacy of the AAV gene therapy for SOD1-ALS and highlights the strong potential for clinical translation of the medium and low-doses. In fact, these doses induce a survival extension of a severe mouse model of ALS-SOD1 and no off-target effects, with the added advantage of requiring only a single injection.

## Materials and Methods

### Animals

SOD1^G93A^ mice (B6SJL-Tg (SOD1*G93A)1Gur/J, Jackson, SN 2726), were purchased from Jackson Laboratory. The animals were housed in a temperature and humidity-controlled room (22°C, 55% humidity) on a 12/12h light/dark cycle, with access to food and water *ad libitum.* Housing conditions adhered to the European guidelines outlined in Directive 2010/63/UE for the care and use of experimental animals. All procedures were approved by the French Ministry of Higher Education and Research and humane endpoints established in the animal protocol (Agreement number #19912). The colony was maintained according to the guidelines provided in the « Working with ALS Mice » manual (https://resources.jax.org/guides/guide-working-with-als-mice). Experiments were performed on sex-balanced groups to account for gender differences in disease progression.

Genotyping of SOD1^G93A^ mice was carried out by PCR and hSOD1 genome copy number assessed as described by Jackson Laboratory.

### AAV Productions

The scAAVrh10-U7-hSOD1 vectors were produced at a single use bioreactor scale (200L) by plasmid triple transfection of Genethon’s proprietary HEK293T cell line using PTG1plus transfection reagent. After 3 days of production, vectors were harvested and lysed with Triton X-100 before clarification. The clarified bulk was loaded in affinity chromatography and the eluate filtrated before a concentration/filtration step. The concentrate was then filtered in 0.2µm filter to obtain the final product.

### AAV Injections

SOD1^G93A^ mice at 50 days of age were administered the scAAVrh10-U7-hSOD1 viral preparation by intracerebroventricular (i.c.v.) injection under anesthesia performed by an intraperitoneal (i.p.) injection of a ketamine/xylazine (100 mg/kg ketamine, Imalgene, Merial, and 10 mg/kg xylazine, Rompun 2%, Bayer; 0.1 mL per 20 g of body weight). Mice were stereotactically injected in the right lateral ventricle (coordinates: 0.2 mm anterior-posterior, ± 1 mm medio-lateral, and 1.8 mm dorso-ventral from bregma) at three different doses: 6 x 10^13^vg/kg, 1.2 x 10^14^vg/kg or 2.4 x 10^14^vg/kg (1µl per gram of body weight administered at a rate of 1.5µl/min).

The comparison among routes of administration was performed on 50-day-old SOD1^G93A^ mice injected with scAAVrh10-U7-hSOD1 that was produced as research-grade vector in our laboratory, as previously described (25).

### Virus vector genome copy number analysis

Total DNA was extracted from freshly frozen tissues using the MagNA Pure 96 DNA and viral NA small volume kit (Roche Diagnosis, Basel, Switzerland) according to manufacturer’s instructions. VGCN were quantified by qPCR using specific primers for the ITRg and normalized to the titin gene copies measured in each sample. qPCR was performed with the StepOne Plus Real Time PCR System (Applied Biosystems) using SYBR Green Master Mix (Thermo Fisher Scientific) under the following thermal cycling conditions: 10 min at 95°C, followed by 40 cycles of 15 s at 95°C and 1 min at 60°C.

The following primers and probes were used:

ITRG forward: 5′-CTCCATCACTAGGGGTTCCTTG −3′,

ITRG reverse: 5′-GTAGATAAGTAGCATGGC −3′,

Probe: 5′-TAGTTAATGATTAACCC −3′

Titin forward: 5′-TTCAGTCATGCTGCTAGCGC-3′,

Titin reverse: 5′-AAAACGAGCAGTGACGTGAGC −3′,

Probe: 5′-TGCACGGAAGCGTCTCGTCTCAGTC-3′.

### Behavioral Analysis and actimetry

Body weight, strength, motor coordination, and neuroscore were assessed twice a week. Grip strength was measured using a grip strength meter (Bioseb). Peak force (in grams) was measured by pulling the mouse back from a metal grid, recorded five times per animals. Motor coordination was evaluated using an accelerating Rotarod (Bioseb), starting at 4 rpm/min. The time and speed spent on the rotarod before the fall of the mice were recorded two times per animal. Neuroscore was assessed based on hindlimb extension evaluation (0 : pre-symptomatic; 1 : first symptoms; 2 : onset of paresis; 3: paralysis or minimal joint movement; 4: humane endpoint) according to published guidelines (71).

The Regularity disturbance index (RDI) and Animal locomotion index were monitored continuously using the Digital Ventilated Cage system (DVC, Tecniplast), which allows automated 24/7 data collection from the home cage. Data were compiled weekly and analyzed by daytime, nighttime and total time.

### Echocardiography

At 105 days of age, mice were lightly anesthetized with 0.5% isoflurane, 2L/min oxygen flow rate and placed on a heating pad (37°C). Echocardiography was performed using 9-14 MHz ultrasound probe (Vivid7 PRO apparatus; GE Medical System Co).

Two-dimensional guided Time Motion mode was used to record left ventricular (LV) measurements from a parasternal long-axis view, including end-diastolic and end-systolic interventricular septum thickness (IVSd and IVSs), posterior wall thickness (PWd and PWs), internal diameter (LVEDD and LVESD), and heart rate (HR). At least three sets of measurements were recorded from three different cardiac cycles. Fractional shortening (FS) was calculated using the formula: [(LVEDD − LVESD)/LVEDD] × 100 and h/r: [left ventricle diastolic wall thickness / radius].

### Hematology and serum bioanalysis

Hematologic parameters were measured using the Vet abc Plus+ (scil). Blood was collected in EDTA tubes from the submandibular vein of the mice using a lancet (Dutscher), and 10 µL of blood were used for analysis.

For serum analysis, blood was collected from the submandibular vein, and serum obtained after 10 min at centrifugation 2000g. Ten microliters of undiluted serum were deposited on Fuji dri-chem slide and concentrations assessed with the FUJI DRI-CHEM NX500 machine. The following biomarkers were measured: aspartate aminotransferase (ASAT), alanine aminotransferase (ALAT), creatinine (CRE), alkalin phosphatase (ALP), creatinine phosphokinase (CPK) and blood urea nitrogen (BUN).

### Regulatory T cells staining by Fluorescence-Activated Cell Sorting (FACS)

Lymph nodes and spleen were collected in RPMI 1640 medium and filtered in a 0.40 µM cell strainer. Lymphocytes were isolated using Ficoll (density 1.077, spin 20 min at 2200 g) and resuspended in RPMI. For staining, cells were incubated with the following antibodies: anti-CD3 (145-2C11, PerCP-Cyanine5.5, 1:20; #45-0031-82, ThermoFisher Scientific), anti-CD45 (FITC, 1:50; #BLE103107, Ozyme), anti-CD8a (AF700, 1:50; #BLE100730, Ozyme), anti-CD4 (Pac Blue, 1:50; #BLE100531, Ozyme), anti-CD25 (PC61.5, eFluor 506, 1:50; #69-0251-82, eBioscience), diluted in PBS 0.5% BSA. Cell viability was determined using LIVE/DEAD Zombie Yellow (1:100, #BLE423104, Ozyme) and was followed by fixation and permeabilization with the BD Cytofix/Cytoperm kit (#AB_2869008, BD Biosciences) for FOXP3 staining (FJK-16s, PE, 1:20; #12-5773-82, eBioscience). Multi-colored immunofluorescence was analyzed on the CytoFLEX (Beckman Coulter) flow-cytometer using Cytexpert software. Data were quantified using FlowJo software (Tree Star).

### Cell culture

Human iPSC from a SOD1^N139D^ ALS patient were differentiated into MN within 3 weeks, as previously described (32). Briefly, iPSC cultured as Embryoid Bodies (EBs) were differentiated into MN progenitors within 10 days. These EBs were then dissociated into single cells and plated on coverslips pre-coated with polyethyleneimine (2mg/ml, Sigma) and laminin (20 µg/mL, Sigma). MN progenitor differentiation into MN was obtained within 4 days of culture in Neuronal Basal Medium (NBM) composed of: 1:1 DMEM/F-12/Neurobasal™, supplemented with N2, B27™ (without vitamin A Thermo Fisher Scientific), L-ascorbic acid (0.01 μM, Sigma), retinoic acid (0.1 µM, Sigma), SAG smoothened ligand (0.5 µM, Enzo Life Sciences), BDNF, GDNF (10 ng/mL, Miltenyi Biotec), and DAPT (10 µM, Sigma). Then, generated MN were incubated in NBM with DAPT, BDNF and GNFF for 3 more days, and then kept in NBM supplemented with neurotrophic factors until the end of the experiment. The medium was changed every other day by replacing only half of the medium.

Primary dermal fibroblasts from SOD1-ALS patients carrying the G93D (Patient 1) and L144F (Patient 2) mutations were provided by the Eurobiobank (Dr. A. Ratti) and were immortalized using established protocols (72) by the Myoline facility (Paris, France) and cultured in DMEM with 10% FBS. Fibroblasts were transdifferentiated into myoblasts using a MyoD transduction protocol (32).

### Viral transduction

MN progenitors were plated at 3×10^5^ cells/well in 24-well plates. Cells were transduced with 10^6^ MOI of scAAVrh10-U7-hSOD1 or scAAVrh10-U7-CTL after 7 days of maturation. At 24 days of differentiation MNs were fixed with 2% formaldehyde for immunofluorescence or, after a wash in phosphate-buffered saline (PBS), stored at −80°C for RNA analysis.

Myoblasts were plated at 2.4 x10^6^ cells in 10mm dishes for RNA analysis. Cells were transduced the day after with 5 x10^6^ MOI of scAAVrh10-U7-hSOD1 or scAAVrh10-U7-CTL for 72 hours. Cell pellets from the 10mm dishes were obtained by centrifuging cells at 3000 rpm for 5 min at 4°C twice, and stored at −80°C.

### RT-PCR and qPCR Analysis

Total RNA was extracted from frozen tissues using Trizol (Invitrogen) and RNeasy Mini kit extraction (Qiagen). cDNA was synthesized from 500 ng RNA and qPCR was performed using Taqman assays for hSOD1 and titin as an endogenous control. StepOne Plus Real-Time PCR System (Applied Biosystems) was used for amplification, and relative gene expression was calculated using the delta-delta Ct method.

### Western Blot Analysis

Protein lysates were extracted from frozen tissues using RIPA buffer enriched with a protease inhibitor cocktail (Complete Mini, Roche Diagnostics) upon tissue homogenization (BeadBug 6, Benchmark Scientific). Protein lysates were quantified using the DC protein assay kit (Bio-Rad). 10 µg of proteins were separated on a 12% polyacrylamide gel (Criterion XT 12% bis-Tris, Bio-Rad) and transferred to PVDF membranes. The following antibodies were used for detection; anti-α-tubulin (1:10000; #T5168, Sigma-Aldrich), anti-actin (1:1000; #A2066, Sigma-Aldrich), anti-human SOD1 (1:1000; #556360, BD Pharmingen) and anti-mouse SOD1 (1:500; #AB5482, Merck Millipore). Peroxidase-conjugated anti-mouse (1:20000; #10179-130, VWR) and anti-rabbit (1:50000; #10014-116, VWR) immunoglobulin (Ig) were used as secondary antibodies. Western blots were developed using the SuperSignal West Dura kit (ThermoFisher Scientific) in a ChemiDoc Imaging Systems (Bio-Rad), and densitometric analyses were performed using Image Lab software (Bio-Rad).

### ELISA

Cu/ZnSOD Human ELISA Kit (#BMS222, ThermoFisher Scientific) was performed following the manufacturer’s instruction, using serum (1:200 dilution) and cerebrospinal fluid (CSF) (1:60 dilution). For CSF collection, mice were placed on a stereotaxic frame under general anesthesia (i.p. injection of ketamine (200 mg/kg, Imalgene, Merial) and xylazine (20 mg/kg, Rompun 2%, Bayer). The needle of the stereotaxic apparatus was inserted into the cisterna magna and the CSF aspirated.

Mouse Troponin I Type 3 (cardiac) ELISA Kit (Colorimetric) (#NBP3-00456, bio-techne) was performed following the manufacturer’s instruction, using heart protein lysate (1:20 dilution).

Similarly, the mouse D-dimer, D2D ELISA Kit (#CSB-E13584M, Cusabio Biotech) was used for cortex protein lysate (1:500 dilution). All measured concentrations were normalized to total protein concentration.

For AAVrh10 levels assessment, a standard curve (Mouse IgG standard; #I5381, Sigma-Aldrich) was prepared in Carbonate buffer (0,0875M Na2CO3, 0.0125M NaHCO3, pH 9.5) and 50 µL of solution analyzed in duplicate in a 96 wells plate. Remaining wells were coated with 50 µL of AAVrh10 capsids in Carbonate buffer at 5^E^8 vg/well and incubated overnight at 4°C. Plates were then washed with PBS + 0.5% Tween20 and blocked in PBS + 0.05% Tween 20 + 2% BSA 2h at room temperature. After washing, 50 µL of blocking solution were added to standard wells and 50 µL of serum (1:4000) in corresponding wells for 1h at RT. Plates were washed 3 times and 100 µL of HRP-conjugated antibody (1:40000; #AP308P, Sigma-Aldrich) in blocking buffer added to each wells and incubated 1h at RT. After washing, plates were incubated 10min with 50 µL of TMB solution (Sigma-Aldrich) and reaction stopped with H2SO4 (3M). Absorbance was read at 450 nm on a TECAN M200 microplate reader.

### Multiplex assay

Several cytokines - Il1β, IL6, IL10, IL12p70, TNFα, IL2, MICP-1 and IFNγ - were simultaneously quantified in brain cortex, spinal cord and liver protein lysates using the Milliplex Map Mouse High Sensitivity T Cell Panel Assay (#MHSTCMAG-70K, Merck Millipore) according to the manufacturer’s instructions. All measured concentrations were normalized to total protein concentration. Samples were processed in duplicate and read on the Bio-Plex MAGPIX Multiplex Reader using Bio-Plex Manager MP software. Data were analyzed using the Bio-Plex Manager 6.1. Fluorescence intensity was calculated with a five-parameter logistic equation (5PL).

### Histological Analyses and Microscopy

Tissues were collected from 70- or 110-days old animals after anesthesia (i.p. injection of ketamine 100 mg/kg and xylazine 10 mg/kg) and double transcardiac perfusion with PBS and 4% paraformaldehyde (PFA, Sigma-Aldrich) in PBS. The spinal cord, dorsal root ganglion (DRG), brain, heart and liver were isolated and stored in 4% PFA at +4°C for 24h, then transferred into a PBS-sucrose solution (30% for spinal Cord, 15% for brain, heart and liver) and stored for at least 24h at +4°C. For the DRG, the tissue was stored in 80% ethanol and then kept at least 24h at room temperature before embedding. All tissues were embedded in Tissue-Tek OCT (Sakura Finetek), and frozen in cold isopentane (between −45°C and −50°C). DRG were embedded in paraffin. Immunofluorescence was performed on spinal cord and brain cryosections, permeabilized in 0.1% Triton X-100 in PBS, subjected to antigen retrieval with citrate buffer (10 mM citric acid, pH 6) for 10 min at 95°C and non-specific epitopes were blocked in 5% BSA (IgG free, protease free, Jackson Laboratory), 0.1% Triton X-100 and 10% normal goat serum (ThermoFisher Scientific). Sections were then incubated overnight at +4°C with the following antibodies: rabbit anti GFAP (1:250; #Z033401, Dako) or rabbit anti-Iba1 (1:400, #019-19741, Wako). After washing, sections were incubated with the secondary antibody goat anti-rabbit 488 (1:750; #A-11008, ThermoFisher Scientific), combined with DAPI (1:5000; #D9542, Sigma-Aldrich). Slides were mounted with FluoroMount-G Mounting Medium (Interchim) and imaged using a Spinning-Disk confocal (Prime95 Camera).

Hematoxylin & Eosin staining was performed on 5µM paraffin-embedded DRG and on 14µm cryosections of brain, heart and liver and imaged with a Spinning-Disk confocal (Nikon Camera). All histological analyses were performed in a blinded manner.

### RNAseq

Libraries for sequencing were prepared by the IGenSeq platform (Paris, France) using the Stranded mRNA Kit (Illumina) according to the manufacturer’s specifications using 25 ng of input RNA. Samples were sequenced after barcoding using the Illumina NovaSeqX platform with 100-bp paired-end reads.

Library alignment and differential gene expression analysis. Reads were aligned to hg38 and mm10. Differential expression analyses were performed using DESeq2 (84).

### Statistical Analyses

Statistical significance was assessed using Student’s paired t-test, one-way ANOVA, or two-way ANOVA, depending on the experiment. Survival curves were compared using the log-rank Mantel-Cox test. Results were considered significant when the p-value is under 0.05. All statistical tests were performed using GraphPad Prism software (version 9.5). Comparisons of the change in body weight, strength, rotarod activity and neuroscoring over time were evaluated using a mixed-effects model. Significance for the main effects of group, time, and their interaction was evaluated with the ANOVA function using type-III fixed effects chi-square tests. Tukey’s multiple comparisons were performed for significant group differences.

## DATA AVAILABILITY

The datasets used and analyzed during the current study are available from the corresponding author on reasonable request.

## ACKNOWLEDGMENTS

We would like to thank Nathalie Mougenot for cardiac echography measurements, Bernard Gjata for liver histology assessment and the Genethon CMC platform for viral batch preparation, titration and VGCN assessment. We would also like to thank the MyoImage and Myoline platform (Centre de Recherche en Myologie), the Histomics platform (Institut du Cerveau) and the staff of the IGenSeq Platform for their assistance with the RNA-sequencing library quality control and for providing access to their facilities. This work was funded by the AFM and was finalized with the support of the Institut National de la Santé et de la Recherche Médicale (INSERM), Sorbonne University and Association Institut de Myologie (AIM).

This work was supported by the national program Investissements d’avenir ANR-10-IAIHU-0006 and the ANR11-INBS-011-Neuratris (ICV-3C, ICV-iPS).

## AUTHORS CONTRIBUTIONS

S.P. performed the in vivo experiments, analyzed, and interpreted data, generated figures, and wrote the manuscript. H.J., M.T, C.A. and A.P. performed in vivo experiments. D.M. and E.S. performed in vitro experiments. L.J. interpreted the FACS analyses. A.S and C-T.M. produced vectors and contributed to analysis. R.M. analyzed RDI data. G.B performed histological analyses. DN, G.F.S. and B.L. analyzed the biomarkers. R.A. provided the human fibroblasts used in the study to generate myoblasts. D.B. provided the iPSC-derived MN progenitors. P.S. oversaw data acquisition and interpretation, analyzed the RNA-sequencing data, generated figures and wrote the manuscript. M.G.B. secured funding, designed the research study, oversaw data acquisition and interpretation, and wrote the manuscript.

## DECLARATION OF INTERESTS

The authors have declared that there are no conflicts of interest.

---

**Figure S1.**
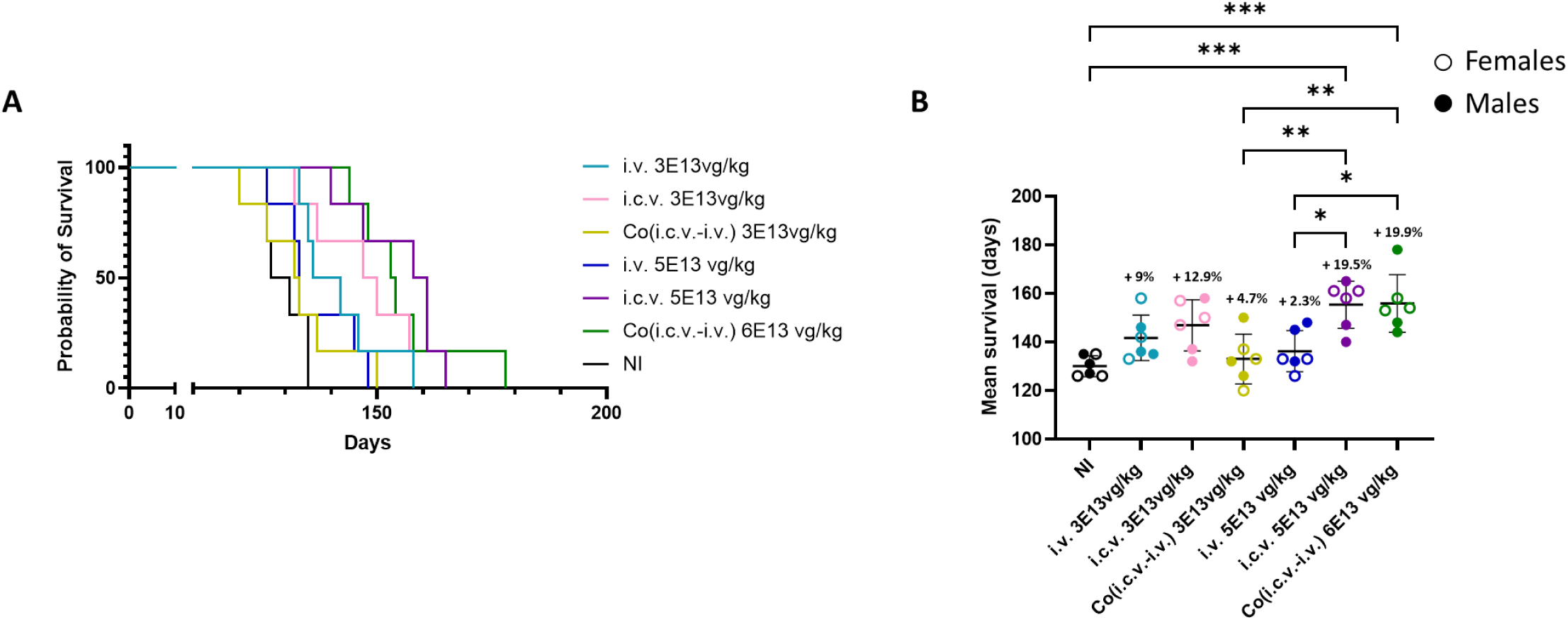
Increased SOD1^G93A^ survival after i.c.v. injection of scAAVrh10-U7-hSOD1 compared to i.v. injection but similar to that after i.v. and i.c.v. co-injection. P50 mice were injected with scAAVrh10-U7-hSOD1 at 3^E^13vg/kg or 5^E^13 vg/kg in the retro-orbital vein (i.v.) or in the lateral ventricle (i.c.v.) or co-injected at 3^E^13vg/kg total (1.5^E^13 vg/kg in i.v. + 1.5^E^13 vg/kg in i.c.v.) and 6^E^13 vg/kg total (3^E^13 vg/kg in i.v. + 3^E^13 vg/kg in i.c.v.). **A.** Kaplan-Meier survival curves of P50 SOD1^G93A^ mice (n=6, sex-balanced groups) injected with scAAVrh10-U7-hSOD1. Differences between the curves were analyzed using the log rank Mantel-Cox test. Statistical differences were observed in the i.v. 3^E^13vg/kg treated group (*p=0.0102), i.c.v. 3^E^13vg/kg treated group (** p=0.0041), and co(i.c.v-i.v) 6^E^13 vg/kg treated group (***p=0.0007) compared to NI. **B.** Scatter plot representation of mean survival of scAAVrh10-U7-hSOD1 injected mice compared to NI mice. Data are expressed as the mean ± SD, and differences between groups were analyzed by one-way ANOVA Tukey’s multiple comparison test. *p < 0.05, ** p < 0.01, ***p < 0.001. All percentages of increased survival on the graph are compared to the mean survival of NI mice.

**Figure S2.**
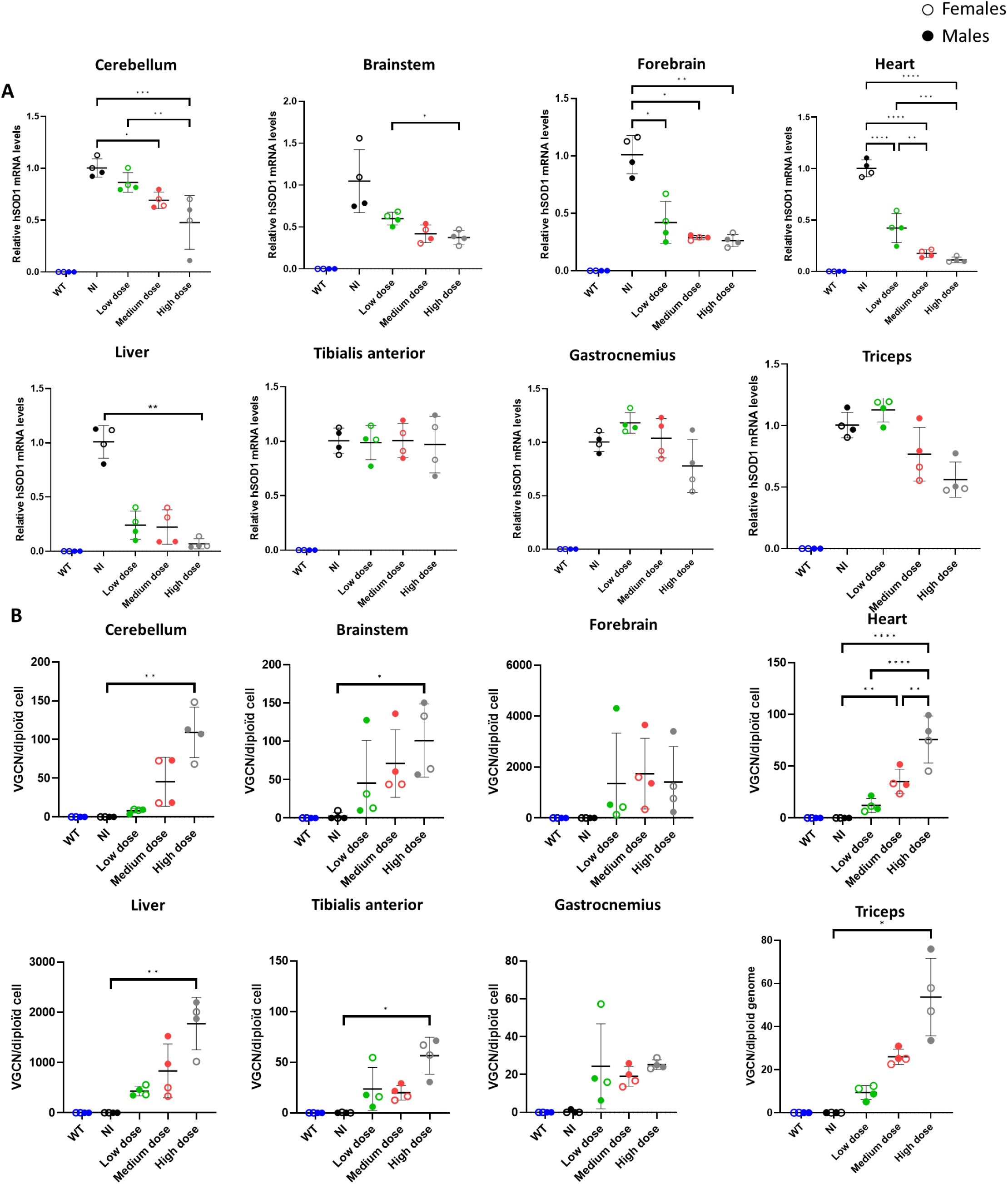
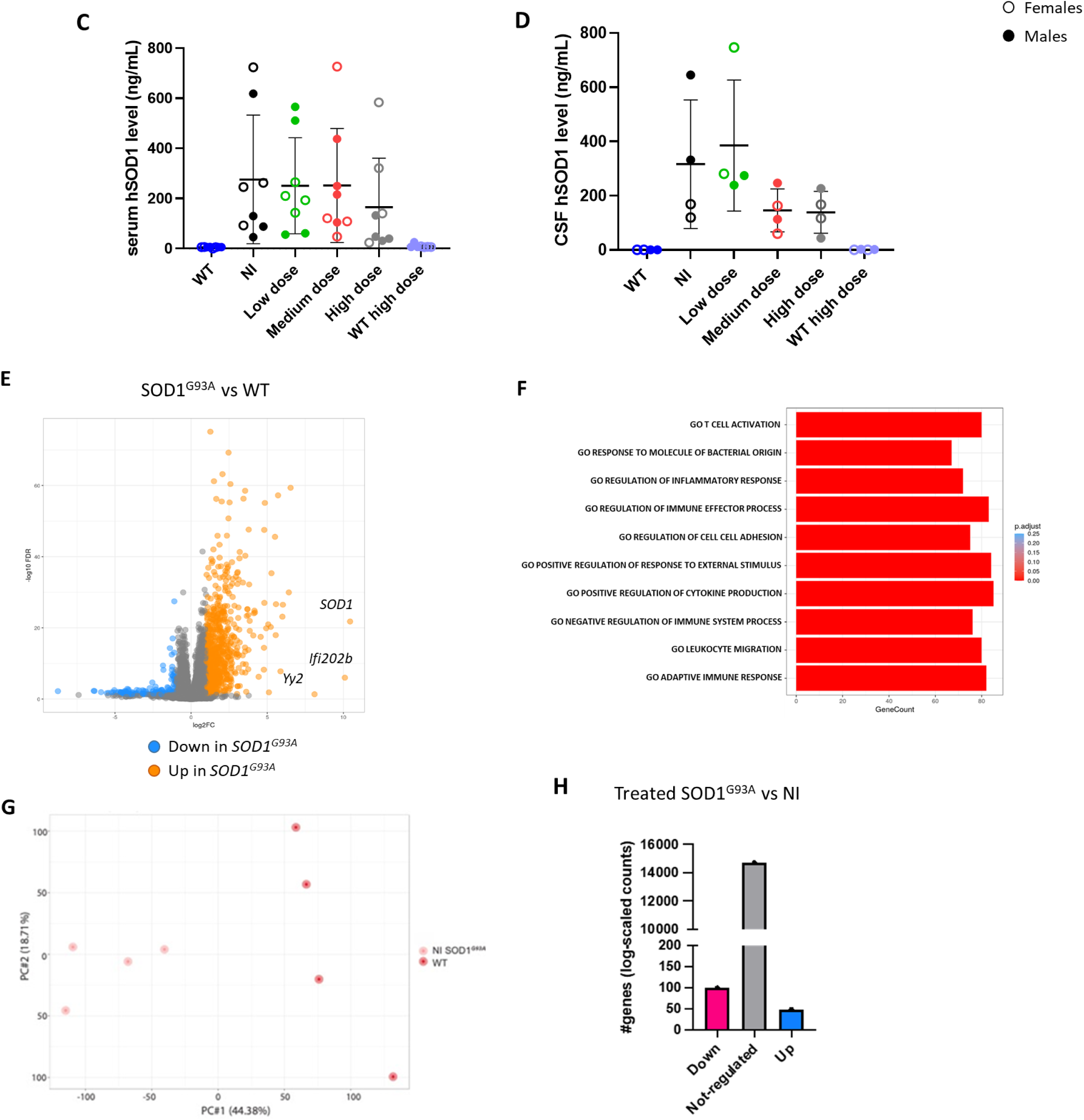
i.c.v. administration of scAAVrh10-U7-hSOD1 results in a dose-dependent decrease of hSOD1 levels in the CNS and the periphery of SOD1^G93A^ mice. **A.** qRT-PCR analysis of *hSOD1* mRNA levels of injected and non-injected mice at P110 (n=4, sex-balanced groups) normalized to *HPRT* mRNA and expressed as relative to levels in NI mice in cerebellum, brainstem, forebrain, heart, liver, tibialis anterior, gastrocnemius and triceps. **B.** Vector genome copy number (VGCN) per diploid cells in cerebellum, brainstem, forebrain, heart, liver, tibialis anterior, gastrocnemius and triceps of injected and non-injected mice at P110 (n=4, sex-balanced groups). **C, D.** hSOD1 levels of injected and non-injected mice at P110 in the serum (n=8, sex-balanced groups) (C) and the cerobrospinal fluid (CSF, n=4, sex-balanced groups) (D). **E.** Volcano plot representing upregulated and downregulated genes in SOD1 vs WT spinal cord extracts; LogFC= 1.5 and p-value=0.05. **F.** Pathway analysis associated with the genes that are differentially expressed in the SOD1^G93A^ spinal cord samples compared to WT. **G.** PCA plot obtained using the variably expressed genes in each spinal cord RNA-seq library of NI SOD1^G93A^ mice vs WT. The variance explained for each principal component (PC1) is plotted on the axis. **H.** Differentially expressed genes from bulk RNA-sequencing analysis of triceps muscle extracts of treated SOD1^G93A^ mice compared to NI at P110 (n=3 for SOD1 group-n=2 males, n=1 female- and n=4 sex-balanced for NI). Data are expressed as the mean ± SD, and differences between groups were analyzed by one-way ANOVA. *p < 0.05, ** p < 0.01, ***p < 0.001, ****p < 0.0001. Significances with WT group are not shown on graph.

**Figure S3.**
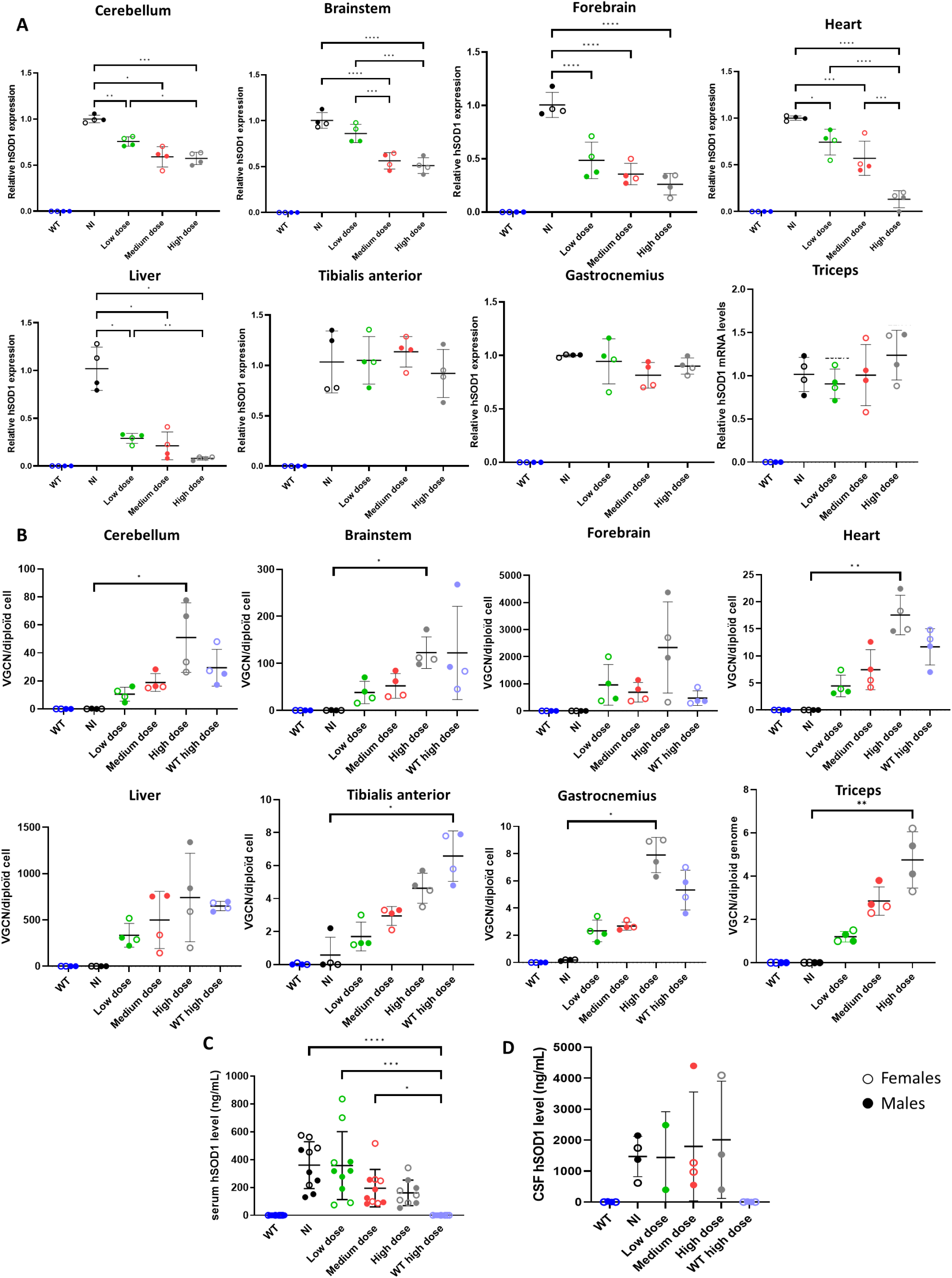
i.c.v administration of scAAVrh10-U7-hSOD1 in SOD1^G93A^ mice induces an efficient and dose-dependent decrease in hSOD1 levels in the CNS and periphery early after injection. **A.** qRT-PCR analysis of hSOD1 mRNA levels of injected and non-injected mice of P70 (n=4, sex-balanced groups) normalized to HPRT mRNA and expressed as relative to levels in NI mice in cerebellum, brainstem, forebrain, heart, liver, tibialis anterior, gastrocnemius and triceps. **B.** Viral genome copy number (VGCN) per diploid cells of injected and non-injected mice of P70 (n=4, sex-balanced groups) in cerebellum, brainstem, forebrain, heart, liver, tibialis anterior, gastrocnemius and triceps. **C, D.** hSOD1 levels of injected and non-injected mice of P70 in the serum (n=10, sex-balanced groups) (C) and the cerebrospinal fluid (CSF, n=4, sex-balanced groups) (D). Data are expressed as the mean ± SD, and differences between groups were analyzed by one-way ANOVA. *p < 0.05, ** p < 0.01, ***p < 0.001, ****p < 0.0001. Significances with WT group are not shown on graph.

**Figure S4.**
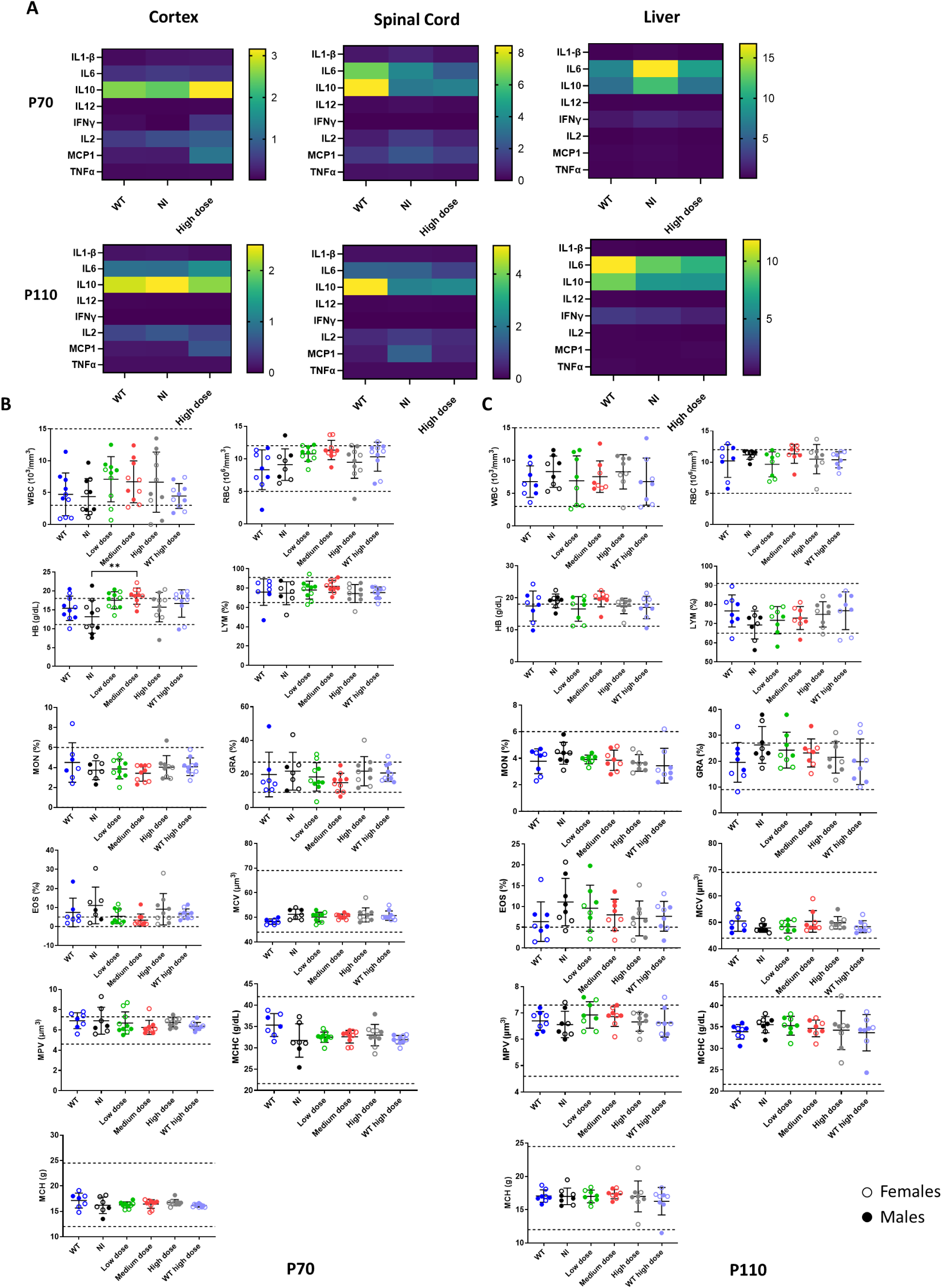
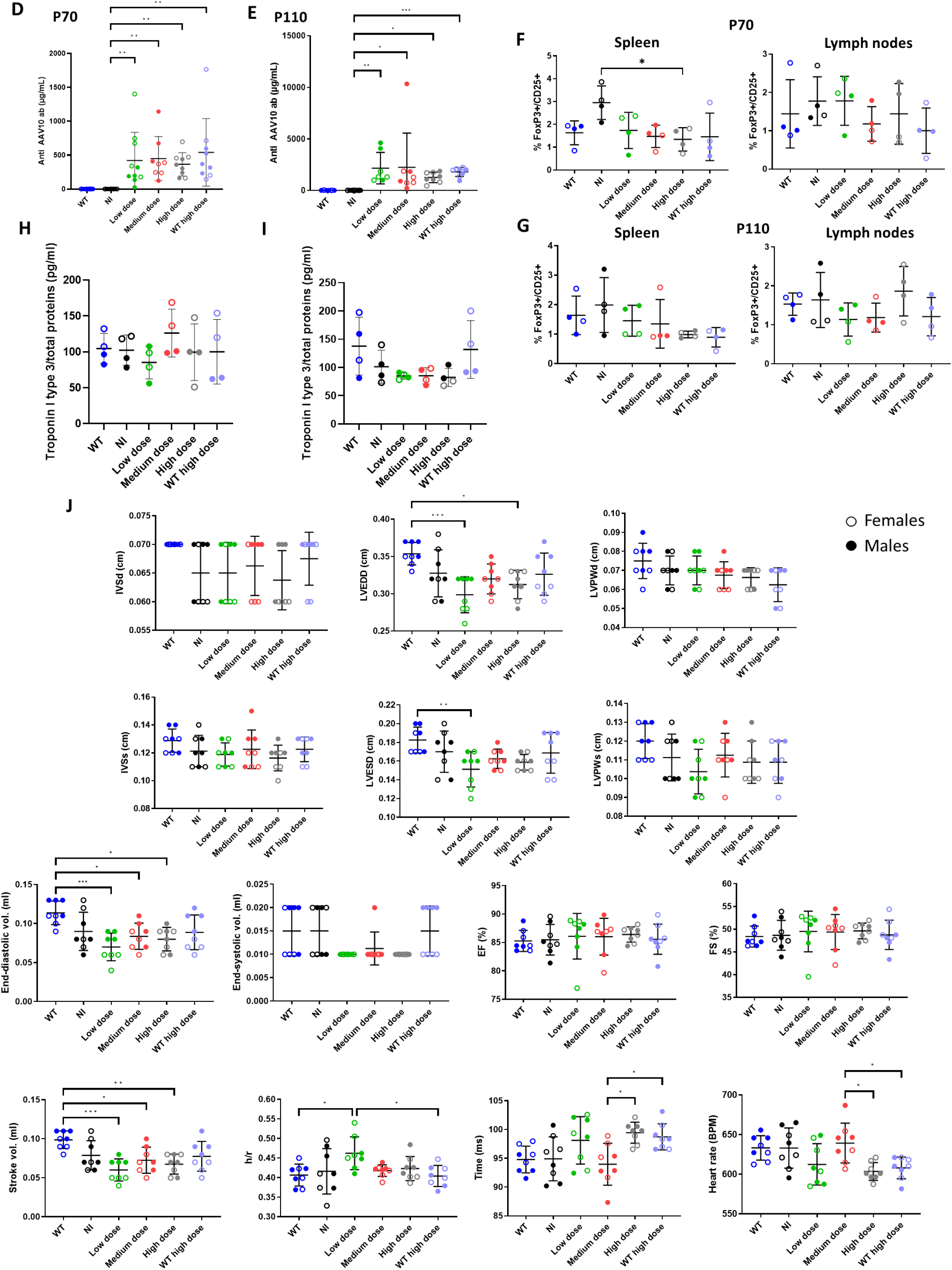
No major immune system activation and heart impairments were observed following i.c.v. scAAVrh10-U7-hSOD1 administration to SOD1^G93A^ mice. **A** Heatmap representation of IL1β, IL6, IL10, IL12, IFNγ, IL2, MCP1 and TNFα concentrations levels in the cortex (left panels), spinal cord (middle panels) and liver (right panels) of injected and non-injected mice (n=4, sex-balanced groups) at P70 (upper panels) and P110 (lower panels). Concentrations are listed in supplementary table 2. **B, C.** Blood parameters analysis of injected and non-injected mice (n=10, sex-balanced groups) at P70 (**B**) and P110 (**C**). The following parameters were assessed: white blood cells (WBC), red blood cells (RBC), hemoglobin (He), lymphocytes (LYM), monocytes (MON), granulocytes (GRA), eosinophils (EOS), mean corpuscular volume (MCV), mean platelet volume (MPV), mean corpuscular hemoglobin concentration (MCHC), mean corpuscular hemoglobin (MCH). Dotted lines correspond to physiological levels. **D, E.** Levels of anti-AAVrh10 antibody (ab) in injected and non-injected mice (n=10, sex-balanced groups) at P70 (**D**) and P110 (**E**). **F, G.** Percentage of regulatory T-cells, corresponding to FoxP3/CD25+ cells, in the spleen (left panel) and the lymph nodes (right panel) at P70 (**F**) and P110 (**G**) of injected and non-injected mice (n=4, sex-balanced groups). **H, I.** Levels of Troponin I type 3 in injected and non-injected mice (n=4, sex-balanced groups) at P70 (**H**) and P110 (**I**). **J.** Cardiac echography parameters analysis in injected and non-injected mice (n=8, sex-balanced groups) at P110. The following parameters were assessed: end-diastolic interventricular septum (IVSd), left ventricular end-diastolic dimension (LVEDD), left ventricular end-diastolic posterior wall dimension (LVPWd), end-systolic interventricular septum (IVSs), left ventricular end-systolic dimension (LVESD), left ventricular end-systolic posterior wall dimension (LVPWs), end-diastolic volume, end-systolic volume, ejection fraction (EF), fractional shortening (FS), stroke volume, h/r, time and heart rate.

**Figure S5.**
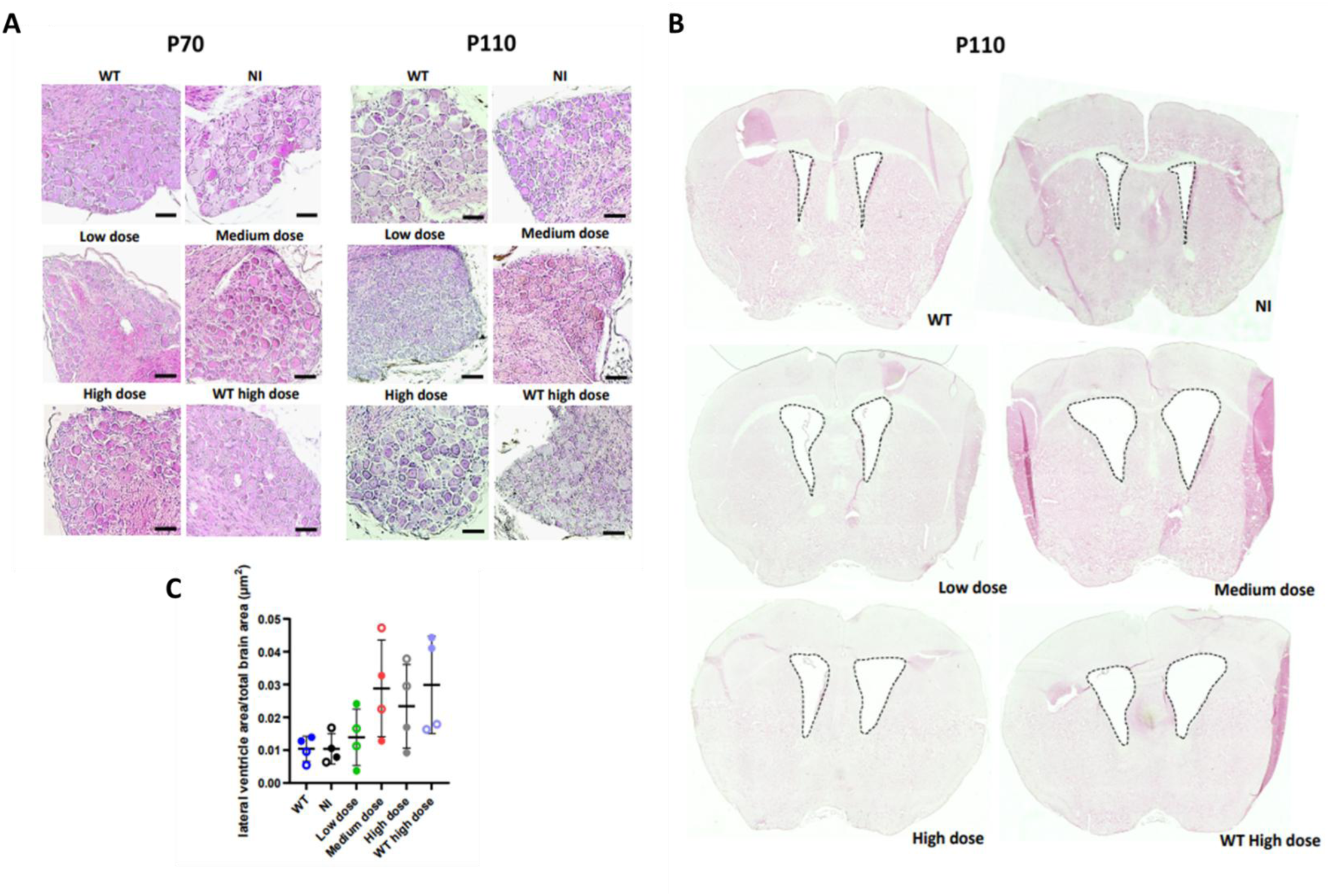
A. No CNS inflammation was observed following scAAVrh10-U7-hSOD1 treatment in SOD1^G93A^ mice but an enlargement of lateral ventricle was noticed in the treated group. **A.** Representative Hematoxylin & Eosin staining of DRG sections from injected and non-injected mice (n=6, sex-balanced groups) at P70 (left panels) and P110 (right panels). Scale bar=50µM. No differences were observed between non-injected and injected mice. **B.** Representative hematoxylin and eosin stain of brain sections from injected and non-injected mice (n=4, sex-balanced groups) at P110 and **C.** Quantification of the lateral ventricle area normalized to total brain area. Compared to control animal, lateral ventricle area (shown in dotted lines on the sections) is increased in injected animal in a dose-dependent manner. Values are expressed as the mean ± SD, and differences between groups were analyzed by one-way ANOVA.

**Supplementary table 1.**
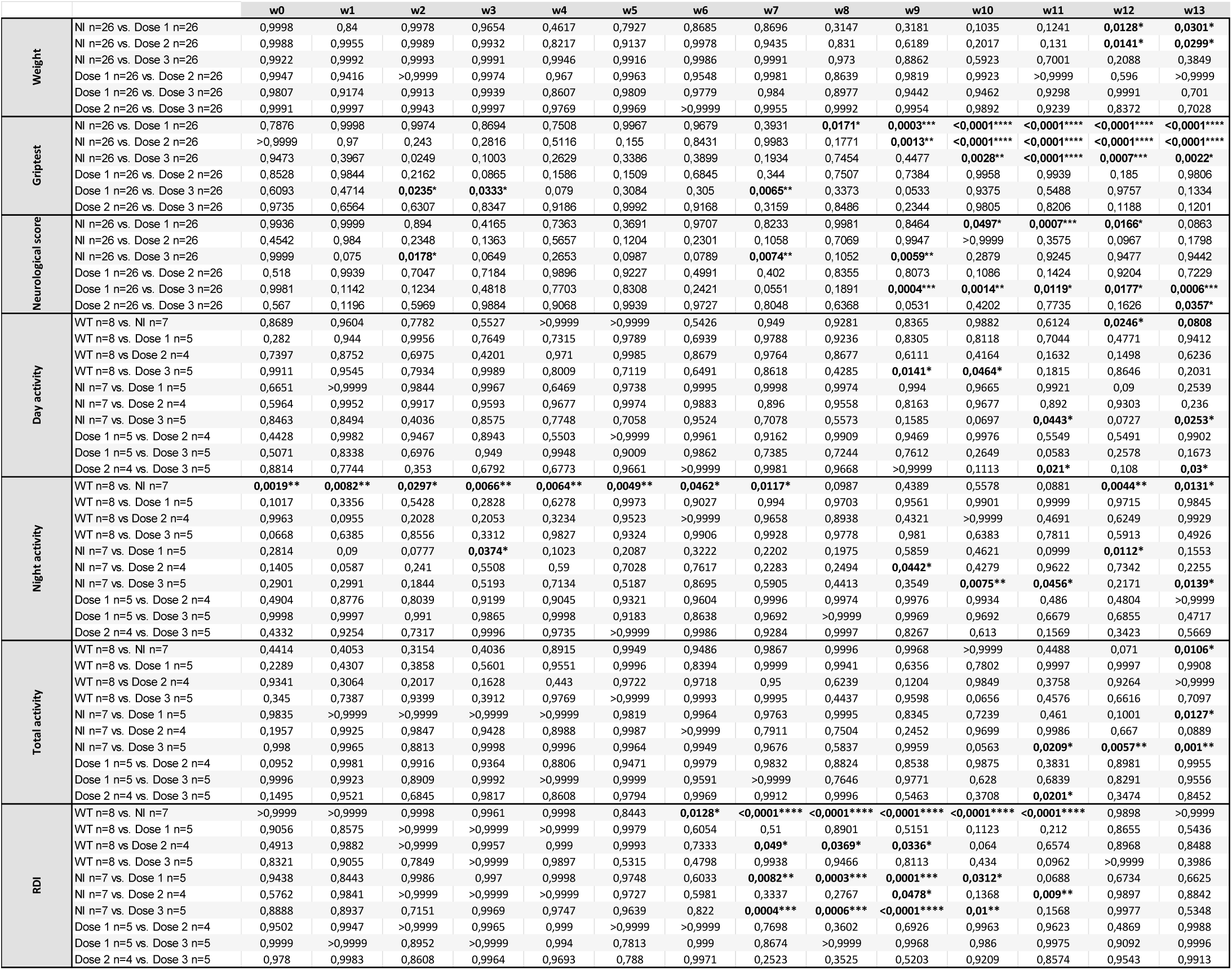

**Supplementary table 2.**
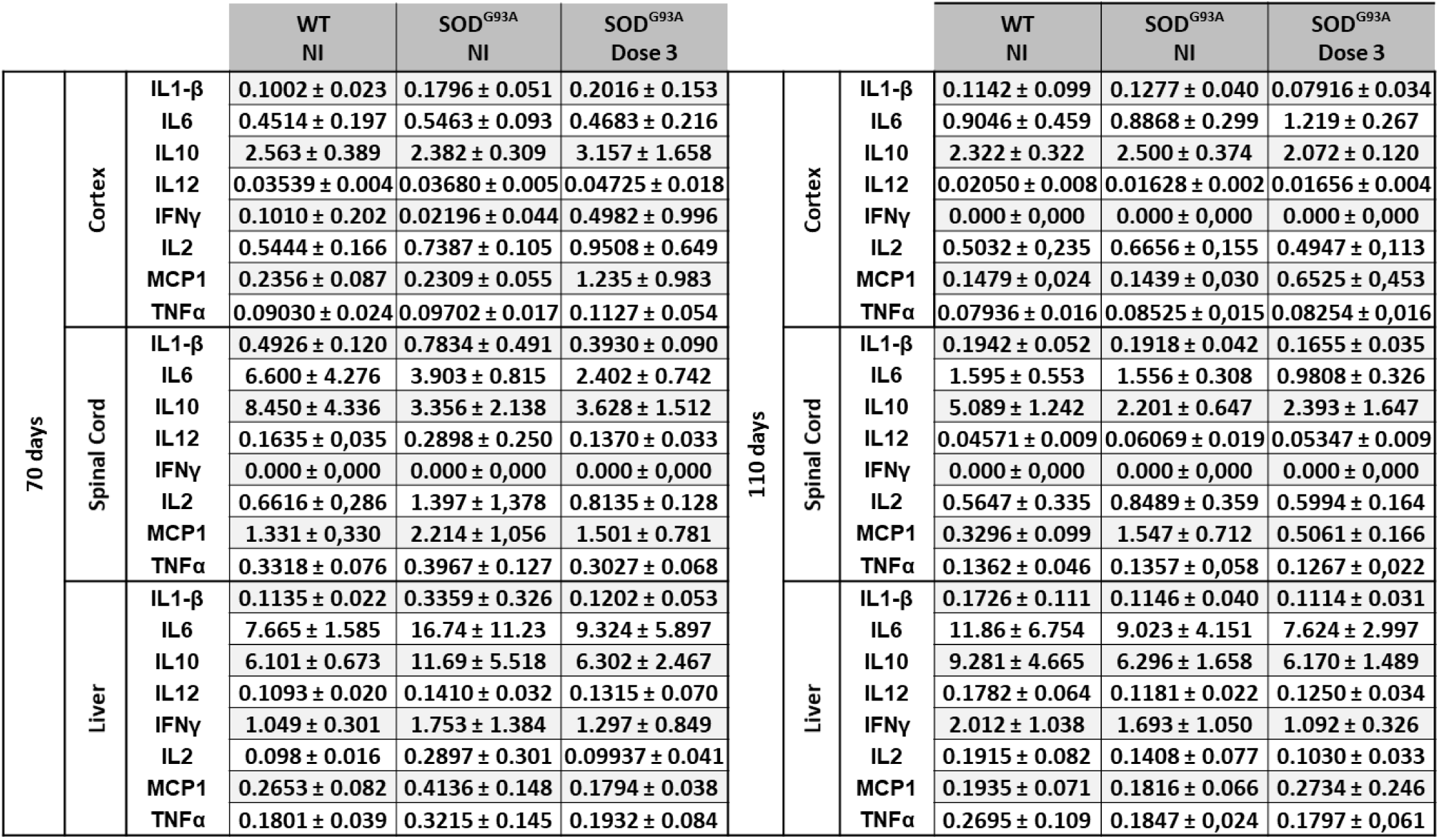

## Notes

### Competing Interest Statement

The authors have declared no competing interest.

